# Accurate detection of metagenomic strain-level associations using average nucleotide identity with StrainSpy

**DOI:** 10.64898/2026.08.30.748153

**Authors:** Sudaraka Mallawaarachchi, Kshitij Tandon, Nikhil Rajan, Vannessa Rossetto Marcelino, Shahneen Sandhu, Sammy Bedoui, Danielle J Ingle, Ashray Gunjur, Gerry Tonkin-Hill

## Abstract

Genetic variation among microbial strains of the same species can profoundly influence their phenotypes, ecological functions, and impacts on human health. Traditionally, the relative abundance of a species has been used to identify associations between the microbiome and disease. However, this approach overlooks intra-species genetic variation and is susceptible to spurious correlations arising from the compositional nature of abundance data and microbial load. Fast, k-mer-based algorithms can now accurately estimate strain-level Average Nucleotide Identity (ANI) in metagenomes. Despite its value as an orthogonal metric for strain-level analysis, existing ANI association methods are limited to case-control designs and rely on arbitrary identity thresholds. To address this, we developed StrainSpy, a statistical algorithm that jointly captures strain variations and replacement across longitudinal and multi-cohort studies without predefined thresholds. In simulations, StrainSpy achieved >98% sensitivity versus 79% for comparable approaches at similar specificity. Analysis of a longitudinal antibiotic study in 12 healthy adults revealed novel strain-level associations across multiple species, including reduced strain-level diversity despite persistence of the species, and showed that a previously reported association with *E. coli* was driven by reduced microbial load. Across 3,414 metagenomes from 17 colorectal cancer cohorts, StrainSpy identified novel strain-level associations in *E. coli* accompanied by enrichment of adhesin genes in addition to recovering well established associations. Importantly, machine learning models built on StrainSpy-selected features matched the predictive accuracy of abundance-based approaches, whilst avoiding the confounding effects of microbial load. StrainSpy is available as an R package at github.com/gtonkinhill/strainspy.

## Introduction

Microbial species exhibit substantial variation in functional and phenotypic capacity as a consequence of genetic differences among individual strains. For instance, *E. coli* strains can differ dramatically in their ecological roles and clinical significance. These range from globally disseminated multidrug-resistant lineages such as the pandemic ST131(1), to strains carrying the polyketide synthase (*pks*) pathogenicity island, which encodes the genotoxin colibactin and has been associated with colorectal cancer mutational signatures(2), to diarrhoeal pathogens including Shiga toxin-producing *E. coli* (STEC) (3) and enteropathogenic *E. coli* (EPEC) (4), to *E. coli* isolated from the secondary environment, wildlife and livestock including poultry and cattle, and commensal *E. coli* (5).

Similar strain-level differences have been observed across many bacterial species within the human microbiota. For example, strains of *Bacteroides* and *Parabacteroides* differ substantially in their capacity to bind and metabolise specific dietary and host-derived glycans(6). Likewise, strain-level variation has been linked to differences in therapeutic responses. For example, bacterial strains have been shown to influence the metabolism of levodopa (L-dopa) in Parkinson’s disease(7). In cancer immunotherapy, only particular strains of *Bifidobacterium bifidum* have been associated with the ability of immune checkpoint inhibitors (ICI) to reduce the tumour burden in mouse models(8).

Despite the established importance of strain-level variation in shaping the structure and function of microbial communities, many tools for microbiome association studies are restricted to the species-or genus-level. Moreover, most current approaches rely on relative abundances to identify microbiome-phenotype associations(9–12). While abundance-based analyses have provided important insights, compositional effects can introduce statistical biases and complicate interpretation(13). Microbial load, for example, is a major determinant of relative abundance variation and is influenced by host factors including age, diet, and medication use(14). There is also ongoing debate regarding best practices for differential abundance testing as different methods frequently yield conflicting results(15, 16).

Containment Average Nucleotide Identity (cANI) is a mea-sure of the genetic similarity between strains present in a metagenomic sample and representative reference genomes. Specifically, cANI quantifies the nucleotide similarity of metagenomic reads that can be attributed to the reference genome, yielding a continuous measure of strain-level similarity. cANI has recently been proposed as an alternative to abundance for microbiome association studies(17). Using cANI as the dependent variable can identify strain-level associations while circumventing many of the methodological challenges associated with the analysis of abundance data. Several fast k-mer based algorithms have been developed for accurately estimating cANI from metagenomes including Sylph(17), sourmash (gather)(18), Mash screen(19) and CMash(20). Despite its value as an orthogonal metric for strain-level microbiome analysis, methods for conducting such association studies remain limited to a narrow range of study designs, such as case-control studies(17). These methods also often require the selection of cANI thresholds below which differences from the reference genomes are ignored, limiting analyses of species that are poorly represented in reference databases.

Alternative approaches for identifying strain-level associations in metagenomics include methods based on phylogenetic regression(21), as well as methods originally developed for genomes from cultured bacterial isolates that have subsequently been applied to binned metagenomic assemblies(22, 23). While these approaches have been successfully applied(24), they typically require an accurate phylogenetic tree to be constructed for each species being considered, which can be challenging in many metagenomic studies. These methods also focus on variation among strains that are detected, rather than jointly modelling strain presence and absence across samples.

To address these challenges, we developed StrainSpy, an algorithm that associates strain-level genetic variation, measured by cANI, with phenotypes across diverse study designs. StrainSpy uses zero-inflated beta regression and phylogeny-aware multiple testing correction to improve its flexibility and statistical power. We demonstrate the accuracy and performance of the algorithm through extensive simulations and application to a well-characterised longitudinal study of antibiotic exposure in 12 healthy adults(25). To further demonstrate the flexibility and scalability of StrainSpy, we reanalysed two separate multi-cohort datasets. The first comprised 3,414 metagenomes from patients with colorectal cancer (CRC), adenoma and healthy controls(24). The second comprised 640 metagenomes obtained from patients receiving immunotherapy for melanoma(26–31) and three types of rare cancers(32). Alongside identifying established and novel strain-level associations in CRC, we showed that StrainSpyderived features readily integrate into standard machine learning frameworks, providing a potential route for disease classification and prediction of treatment response. Across both clinical contexts, StrainSpy performed comparably to conventional abundance-based methods, supporting its use as a robust complementary approach for microbiome analysis. Strain-level features may also yield biomarkers that could be more readily translated into targeted assays, such as qPCR. The StrainSpy R package is available under an MIT licence at https://github.com/gtonkinhill/strainspy.

## Results

### Overview of StrainSpy algorithm

Unlike common abundance based approaches, StrainSpy performs microbiome association tests by modelling cANI, allowing for the analysis of strain-level variation whilst being less sensitive to compositional effects such as microbial load(14). cANI measures the similarity between a reference genome and its closest match in a metagenomic sample. It ranges from 0, indicating absence, to 1, indicating presence of an identical strain. A cANI of 0.95 indicates the presence of a strain with a species-level match in the reference database(17).

As shown in Figure 1, Strainspy uses a zero inflated beta (ZiB) model for association testing to account for both species presence/absence and the underlying sub-species distribution of cANI. By modelling these two components jointly, StrainSpy avoids the need to select a strain-level ANI threshold, addressing a limitation recognised in previous ANI-based association studies(17). An empirical Bayes regularization approach is used to stabilise the model fit and reduce spurious associations, particularly in small datasets. Given the high genomic variability in certain taxa, the exact strain responsible for modulating a phenotype may not be present in the reference database. Importantly, this does not prevent association testing (Figure 1a). When multiple strains of the same species are represented in the database, these references provide a basis for measuring the genetic similarity of observed strains across samples. For example, if strains associated with the phenotype are more common in cases than controls, the resulting cANI distributions relative to those references will differ between groups, even when the causal strain itself is absent from the database (Figure 1a).

**Fig. 1.**
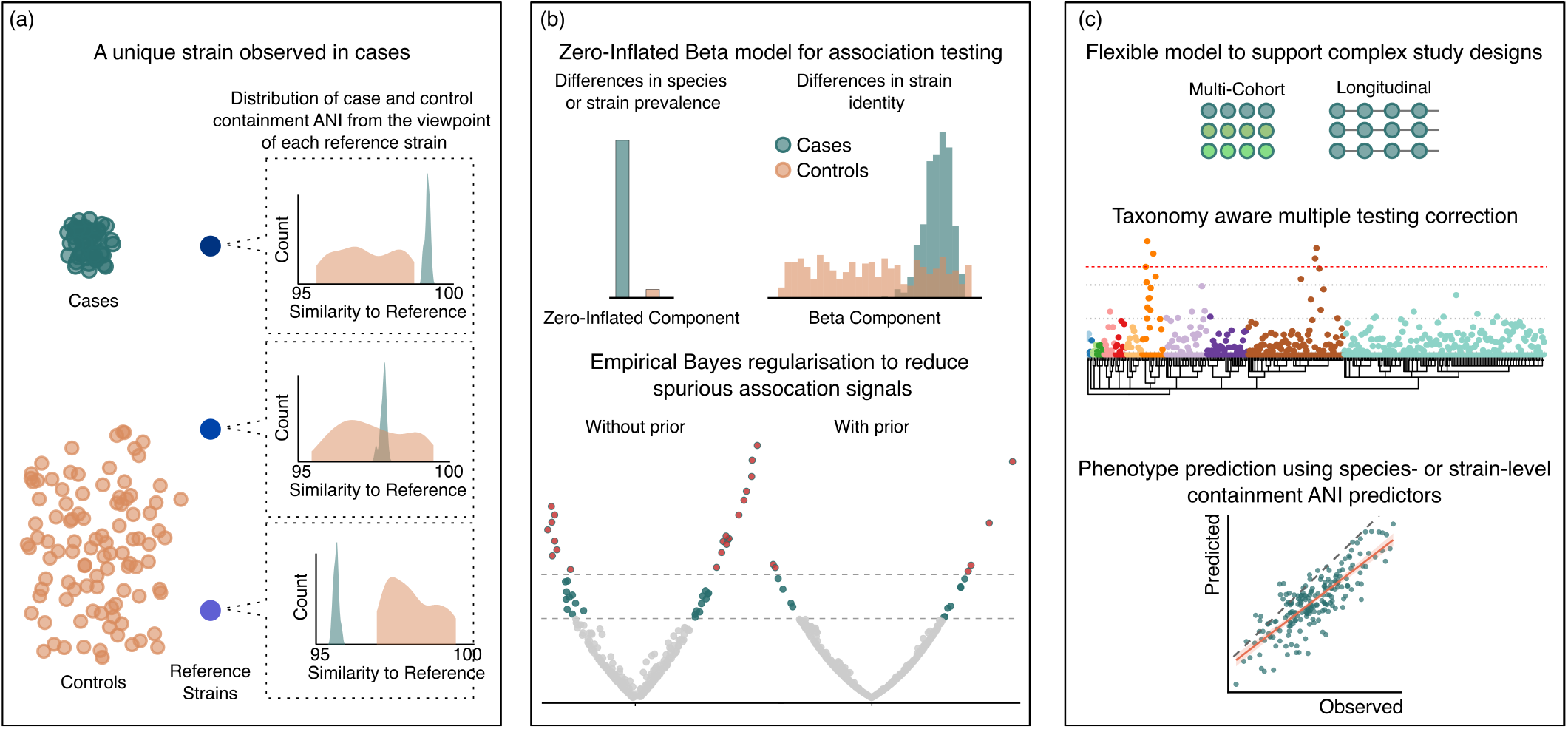
Overview of the StrainSpy workflow. (a): For a given species, each coloured point represents a strain, with distance between points reflecting genetic similarity. Green and orange represent observed strains from each case and control sample, respectively, while the three shades of blue represent reference strains in the database. cANI distributions are shown from the viewpoint of each reference strain by comparing the genetic similarity between each reference and observed strains. Although the causal strain is absent in the database, cases will produce a concentrated cANI distribution compared to controls. (b) Association testing is performed using a zero-inflated beta (ZiB) model, with optional empirical Bayes regularisation to stabilise model fitting and reduce false-positive associations. (c) Containment-ANI can be modelled against diverse predictors to support complex study designs, with phylogenetic-aware multiple testing correction and downstream phenotype prediction from strain-level profiles.

Since Strainspy models cANI as the dependent variable, it is suited to testing for associations across a wide range of complex designs, including multi-cohort, longitudinal, and repeated-measures studies (Figure 1c). Importantly, in addition to standard multiple-testing correction, StrainSpy incorporates a phylogenetic-aware approach using the Harmonic mean p-value(33), increasing the statistical power of studies that consider large numbers of strains. Finally, StrainSpy enables the extraction of strain-level cANI features for downstream predictive modelling using a range of common machine learning approaches.

### StrainSpy reveals strain dynamics following antibiotic exposure

To demonstrate the capability of StrainSpy, we used a well curated longitudinal study of 12 healthy adult males exposed to broad spectrum antibiotics (vancomycin, gentamicin and meropenem)(25). Analysing this type of longitudinal study would have been challenging with previous cANI-based logistic regression approaches(17), which were developed primarily for case-control comparisons and model binary outcomes as a function of cANI predictors.

Faecal samples from each adult were collected at five time points spanning baseline (day 0) and post-exposure (day 4, 8, 42 and 180) following a four-day course of broad antibiotics, used to treat infections caused by multidrug-resistant bacteria. Samples were available from nine individuals on day 4, and from all twelve at the remaining time points.

Consistent with the findings of the original study, StrainSpy identified a substantial reduction in microbial diversity immediately following antibiotic treatment, followed by partial restoration of the community by day 42 and further recovery by day 180(25) (Supplementary Table 1). Relative to baseline, the early post-treatment timepoints (days 4 and 8) showed a widespread reduction in the prevalence of 1,140 species, which is captured by the zero-inflated component of the StrainSpy model. These changes included an increase in the prevalence of the opportunistic pathogen *Klebsiella pneumoniae*, which remained detectable in three subjects at day 180 (Figure 2a). Conversely, a reduction in the prevalence of strains of the short-chain fatty acid (SCFA) producer *Bifidobacterium longum* at day 8 was observed, with recovery in most subjects by day 180. This was accompanied by a transient increase of the oral-associated *Bifidobacterium dentium* (Figure 2a; Supplementary Figure S4) in the gut. Similar increases in oral-to-gut enrichments have previously been associated with altered host health and multiple disease states(34). A similar pattern was observed between *Ruminococcus bromii* and *Ruminococcus gnavus*.

**Fig. 2.**
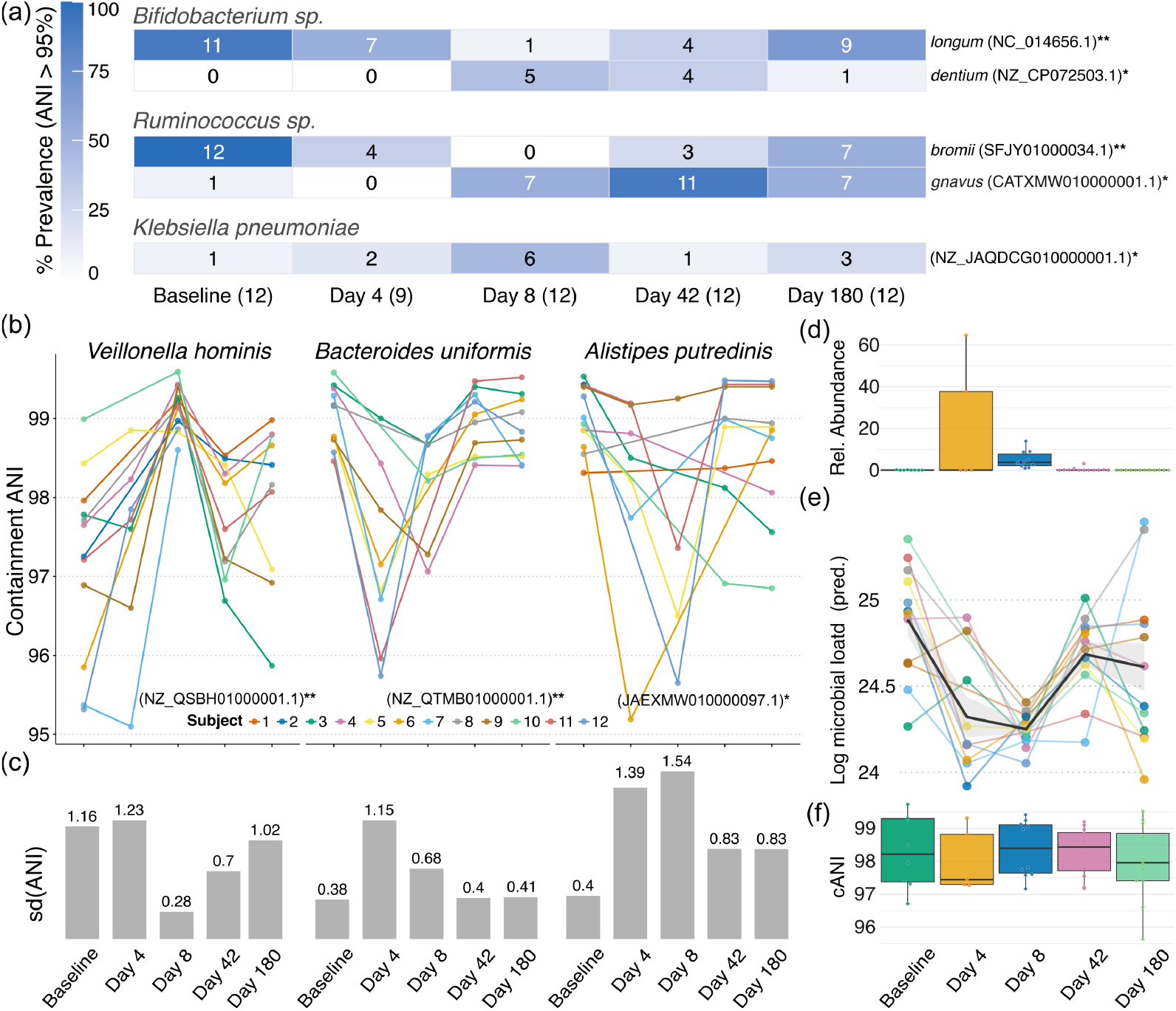
StrainSpy reveals widespread perturbations after antibiotic exposure. (a) Prevalence of significantly associated taxa illustrating strain-level presence, absence, replacement and persistence dynamics. Each tile shows the number of subjects with a detectable strain (cANI > 0). (b,c) Within-subject strain cANI trajectories for selected significant taxa highlight strain turnover events, with standard deviation summarising population-level variability. (d) Relative abundance changes in *E. coli* following exposure produce a significant association under standard abundance-based testing. (e) Predicted microbial load captures overall community depletion at day 8, and inclusion of this covariate removes the apparent association. (f) StrainSpy shows no significant signal for *E. coli* in prevalence (a) or turnover, consistent with reduced sensitivity to microbial load–driven effects.

Crucially, beyond detecting changes in strain prevalence, StrainSpy also identified strain-level cANI shifts within 282 species that persisted despite antibiotic exposure, with 96 associations still present at day 180. For example, *Veillonella hominis* (Figure 2b) exhibited a marked reduction in the standard deviation of cANI values at day 8 (s.d. = 0.28), which recovered to near-baseline levels by day 180 (s.d. = 1.02), despite the species remaining detectable throughout the study. A possible explanation is that antibiotic exposure preferentially depleted more susceptible strains of *V. hominis* while allowing other strains within the species to persist. In contrast, *Bacteroides uniformis* and *Alistipes putredinis* were found to have increased within-species strain diversity following treatment (Figure 2b). For *B. uniformis*, cANI standard deviation returned to near-baseline levels by day 180 (s.d. = 0.38 at baseline; 0.41 at day 180), whereas *A. putredinis* maintained elevated strain divergence (s.d. = 0.40 at baseline; 0.83 at day 180). These patterns are consistent with strain replacement following antimicrobial treatment and suggest strain level restructuring within persistent species that would have been missed by traditional species-level abundance analyses.

Unlike the differential abundance based analysis conducted in the original study(25), StrainSpy did not identify a significant association between *E. coli* cANI and antimicrobial treatment (adjusted p = 0.532), despite the species level relative abundance clearly increasing at day 4 and 8 (Figure 2d). A reanalysis suggests that the original relative abundance based associations were driven by a reduction in overall microbial load following antibiotic exposure (Figure 2e, see methods). Although the association was significant in an abundance-based analysis prior to adjustment (adjusted p = 0.028), it was no longer significant after including predicted microbial load as a covariate (adjusted p = 0.44). This indicates that StrainSpy is robust to abundance-based confounding arising from changes in total microbial load.

### StrainSpy provides robust results in simulated case–control studies

To further validate StrainSpy’s accuracy, we generated simulated case-control studies using an implantation-based framework, adapted from Wirbel *et al*.(16), applied to 200 randomly selected healthy gut metagenomes from Zeevi *et al*.(35) (see Methods). In each simulation we artificially implanted a signal in half the metagenomes by either reducing the cANI distribution of a randomly selected strain or reducing its prevalence by a specified fraction up to 75%. StrainSpy was then run alongside both the logistic regression based approach described in Shaw *et al*.,(17) and an approach based on ordered beta regression which has been shown to accurately model proportional data(36).

All three models compared in Figure 3a detected cANI changes as low as 0.5%. At cANI changes > 1%, the ZIB used by StrainSpy and ordered beta regression models achieved > 98% sensitivity, outperforming logistic regression (~79%) at the cost of a very small drop in specificity (< 0.05%). However, when simulating reduced strain prevalence, the ordered beta model demonstrated poor sensitivity (< 25%) even at strain depletions as high as 75% (Figure 3a). Regardless of the scenario, all models maintained exceptionally high specificity (> 99.99%). Across all simulated changes in strain cANI and prevalence (Supplementary Figure S1), the ZiB model used in StrainSpy consistently yielded higher sensitivity than the logistic regression based approach.

**Fig. 3.**
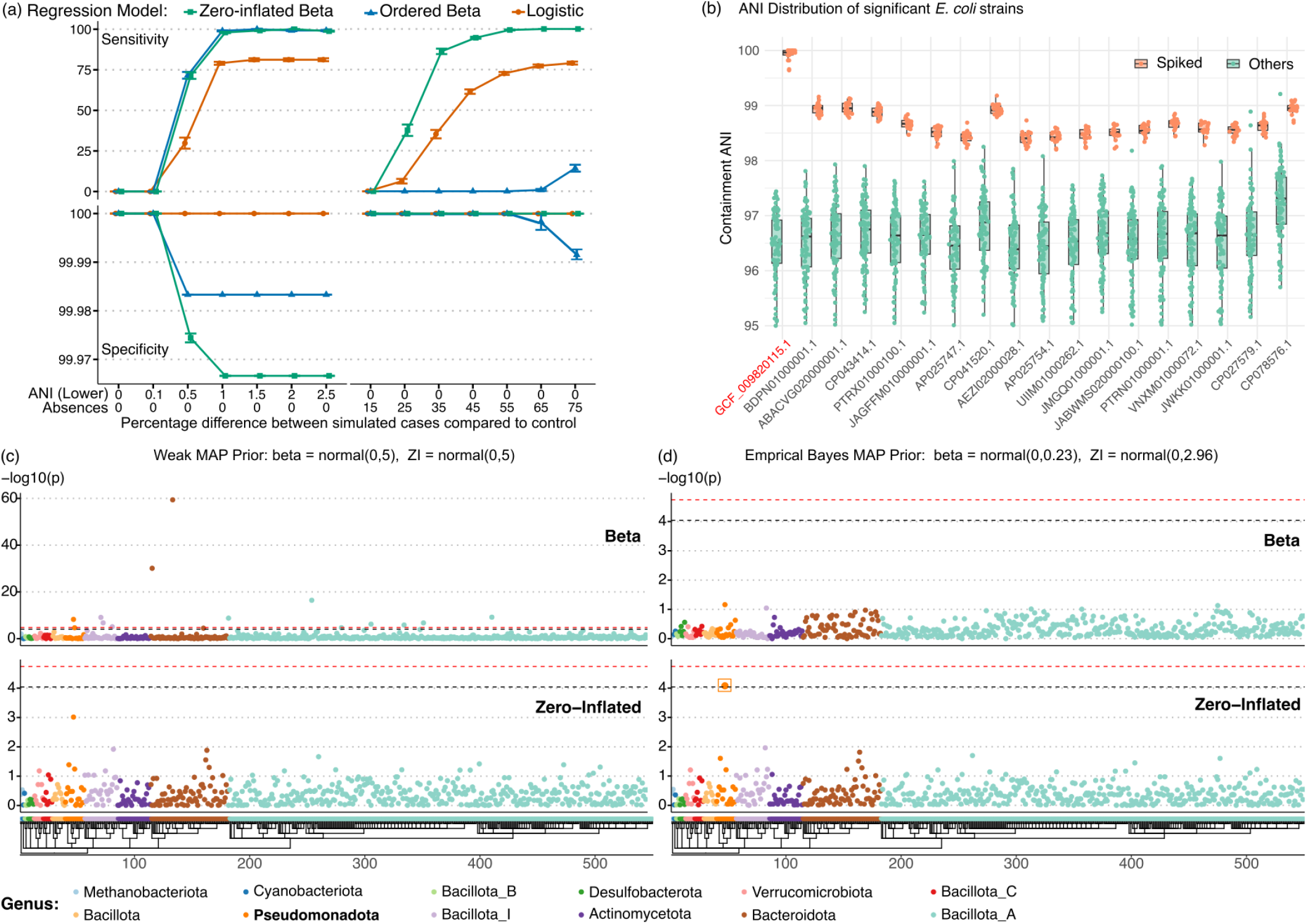
Performance of StrainSpy in simulated case–control and spike-in simulations. (a) Sensitivity and specificity of zero-inflated beta (ZiB), ordered beta, and logistic regression models under simulated strain identity reduction (left) and strain depletion (right). All models detect small ANI changes (0.5%), with ZiB and ordered beta achieving near-perfect sensitivity at >1% ANI change, while logistic regression shows reduced sensitivity. In depletion scenarios, the ordered beta model shows poor sensitivity despite consistently high specificity across all models. (b) Variation in non-zero ANI values for spike-in samples (orange) versus controls (green) across ordered significant hits detected by StrainSpy. Simulated reads from *Escherichia* sp. (GCF_009820115.1) were spiked into orange metagenomes, and the method correctly ranks the spiked genome as the top hit (adjusted p = 1.2 *×* 10^−144^). (c–d) Manhattan plots from ZiB analysis in a small-sample setting (16 cases, 17 controls). The default weak MAP prior results in multiple false positives and failure to detect the true signal (c), whereas an empirical Bayes prior eliminates false positives and recovers the spiked strain from the *pseudomonadota* genus, shown in an orange square (d).

In order to compare to methods that do not make use of cANI we also simulated a strain level signal by spiking in *Escherichia* sp. (GCF_009820115.1) into a subset of the 200 metagenomes. This strain was chosen for spike in as it was sufficiently distant to other *Escherichia* sp. strains detected in the dataset. Using Art(37, 38), short reads were simulated from this genome and a subset of reads were spiked into 20 randomly selected cases maintaining a fold-coverage of 1 *×* (see Methods). StrainSpy identified the spiked-in strain (adjusted p = 1.2 *×* 10^−144^), alongside 17 additional *Escherichia* sp. strains sharing > 97.5% ANI with the spike-in genome (p-values between 1.1 *×* 10^−28^ and 2.38 *×* 10^−74^). These additional strains were co-identified due to their high sequence similarity with the spiked in genome (ANI > 97.5%) and StrainSpy’s assumption of strain independence (Figure 3b; Supplementary Figure S2). To assess the robustness of StrainSpy in a small-sample setting, we repeated the *Escherichia* sp. spike-in analysis with only 16 cases and 17 controls (Figure 3c). Using the weak maximum a posteriori (MAP) prior led to several false-positives and no significant hits were detected in Pseudomonadota (see Methods). How-ever, after using the empirical Bayes MAP prior (see Methods), the spike-in was detected (adjusted p = 1.51 *×* 10^−7^) and no false positives were found.

Next, we attempted to compare these results with AnPan(21) which relies on a user supplied species phylogeny, commonly generated by applying the StrainPhlAn(39) pipeline. Although the species genome bin containing the spike-in strain (SGB10068), was detected in 52.2% of samples, most detections occurred at low abundance (> 90% below 0.25% relative abundance; median relative abundance among detected samples = 0.032%). Consequently, there was insufficient metagenomic coverage for reliable phylogenetic reconstruction using StrainPhlAn(39) preventing a direct comparison with An-Pan.

To enable comparison using a species with sufficient phylogenetic resolution, we instead selected *Parabacteroides distasonis* as the species that had the highest prevalence (> 98%) in the dataset. *P. distasonis* strain GCF_024791025.1 was selected to implant into 20 randomly selected samples, and simulated reads were spiked in at coverages of 1*×*, 3*×* and 5 *×* (see Methods). StrainSpy detected the implanted strain at all coverage levels.

StrainPhlAn(39) detected *P. distasonis* (SGB1934) in 84 samples at sufficiently high coverage to construct a species phylogeny. In contrast to StrainSpy, the StrainPhlAn-AnPan approach did not identify a statistical association when the coverage was only 1 *×* although it did correctly identify associations at 3 *×* and 5 *×* coverage (Supplementary Figure S3). Overall, these results suggest that while AnPan performs well when accurate species-level phylogenies can be constructed, cANI-based approaches are better suited to detecting associations involving rarer strains with limited sample representation or sequencing coverage.

### StrainSpy reveals novel strain-level associations across large multi-study cancer microbiome cohorts

To demonstrate the flexibility and scalability of StrainSpy for large-scale meta-analyses across diverse microbiome cohorts, we reanalysed 3,414 publicly available fecal metagenomes from a recently published meta-analysis of the microbiome in CRC(24). This cohort spanned 17 independent datasets across 10 countries and comprised 1,322 CRC and 614 adenoma cases, and 1,478 controls, covering a broad range of tumour stages and locations. Importantly, because batch effects are difficult to account for in abundance-based microbiome studies, the original study relied on a meta-analysis of standardised mean differences (SMDs) calculated separately within each dataset, rather than estimating effect sizes from batch-corrected abundance data. In contrast, StrainSpy substantially simplified the analysis by allowing all datasets to be jointly analysed in a single model. Study was modeled as a random effect to account for cohort-level variation while estimating strain-level associations across the pooled dataset.

In general, the StrainSpy analysis was concordant with original abundance based findings identifying 64 species with differential prevalence patterns and 9 strain-level associations (Supplementary Table 2).This included that the microbiome in adenoma samples did not differ significantly from controls, suggesting that microbial differences may become more pronounced during later stages of the adenoma-carcinoma transition. In contrast, comparisons between CRC and controls revealed distinct microbiome shifts. The CRC population was characterised by the depletion of several commensal taxa and enrichment of opportunistic and oral-associated taxa (Figure 4a). In particular, SCFA producing taxa such as *Faecalibacterium prausnitzii* and *Butyribacter intestini* were depleted in CRC whereas taxa more commonly associated with oral or opportunistic niches, including *Fusobacterium animalis* (also known as *Fusobacterium nucleatum* subsp. *animalis*, FNA), *Dialister pneumosintes, Peptostreptococcus stomatis*, and *Parvimonas micra*, were more prevalent in CRC.

**Fig. 4.**
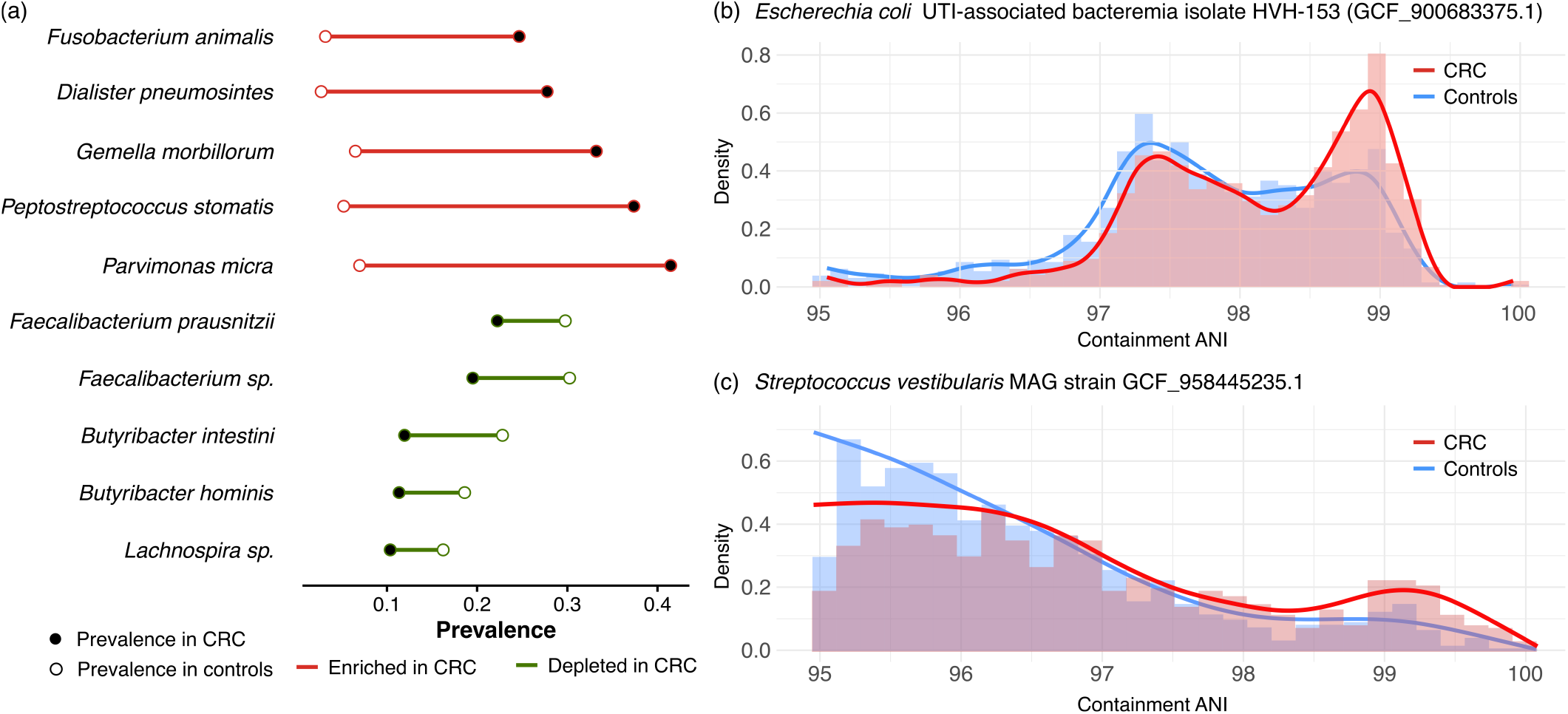
StrainSpy detects microbiome changes in colorectal cancer (CRC) in a large multi-cohort dataset. (a) As reported previously(24), a subset of significant changes in prevalence shown here highlight the enrichment of opportunistic and oral-associated taxa and depletion of SCFA producers in CRC compared with controls. Differences in cANI distribution indicates strain identity differences in (b) *E. coli* and (c) the oral associated species *Streptococcus vestibularis* between CRC and controls.

In contrast to the original meta-analysis, StrainSpy detected a significant shift in the cANI distribution of strains of *E. coli* in CRC (Figure 4b). This included a hit to the Urinary Tract Infection (UTI) associated reference genome GCF_000458275.1 (adjusted p = 0.04), originally sampled from a bloodstream infection. Here, the 0.07% ANI difference corresponds to ~3 − 4 kbp of genome-wide sequence divergence. While the original meta-analysis using AnPan did not capture any *E. coli* strain-level differences, a separate gene level analysis did identify altered *E. coli* gene content in CRC, supporting our findings.

To expand on this previous gene level analysis we investigated whether the presence of *E. Coli* adhesin genes differed significantly between CRC patients and controls as adhesins, both fimbrial and non-fimbrial, have been established as critical for *E. coli* to attach to host epithelial cells. Several adhesin systems, including fimbriae, curli, and autotransporters, were more prevalent in the CRC samples (Supplementary Table 3), including those known to promote attachment to epithelial cells. The *fim* operon *fimABCDEFGHI* encodes type 1 fimbriae that facilitate colonisation and persistence within the host and are an important virulence factor in extraintestinal *E. coli* (ExPEC), particularly uropathogenic *E. coli* (UPEC) (40), where sequence variation in the *fimH* gene is used to provide subgroup resolution within important ExPEC clones (41, 42). Similarly, we observed enrichment of PixB, a homologous regulatory protein to PapB that is the transcriptional regulator of P fimbriae, which were first identified in UPEC and promote adhesion to host epithelial cells (40). Adhesin genes associated with biofilm formation were also identified, including *csgD*(43). Collectively, these findings suggest that the strain-level differences identified by StrainSpy are accompanied by differences in *E. coli* factors involved in adhesion, colonisation and persistence. The intestinal environment of CRC patients is substantially perturbed by the disease and its treatment, such as exposure to antibiotics and chemotherapy and surgical interventions. Such perturbations may impose selective pressures on resident *E. coli* populations, potentially favouring lineages with enhanced capacity for adhesion, colonisation, and potential pathogenesis, in addition to biofilm formation which creates an extracellular matrix that facilitates evasion of antibiotics.

In addition to *E. coli*, StrainSpy also revealed novel strain-level cANI differences in *Streptococcus vestibularis* associated with CRC (Figure 4c). As *S. vestibularis* is predominantly an oral species, this tumor-specific strain-level association parallels recent findings for *Fusobacterium nucleatum*, in which the former subspecies *animalis* has emerged as a distinct clade (now recognised as a separate species) with a preference for colonising the tumour niche(44).

Next, to examine a disease with a weaker microbiome association we evaluated whether a similar multi-cohort meta-analysis of the relationship between the gut microbiome and immunotherapy efficacy could identify strain-level associations. We applied StrainSpy to a combined cohort of publicly available faecal shotgun metagenomes from patients receiving ICI, including combined immune checkpoint blockade (CICB) targeting programmed cell death protein 1 (PD-1) and cytotoxic T-lymphocyte-associated protein 4 (CTLA-4). The cohort comprised three rare cancer datasets (UGB: upper gastrointestinal and biliary cancers, NEN: neuroendocrine neoplasms and GYN: rare gynecological tumors) from Gunjur *et al*.(32), together with six melanoma datasets(26–31), filtered according to the criteria described by Gunjur *et al*.

Similar to previous abundance-based meta-analyses(28), StrainSpy did not identify significant strain-level associations across the combined cohort or within the melanoma-only subset. This remained the case after adjusting for treatment regimen (CICB versus PD-1 monotherapy) as a covariate. However, StrainSpy did identify strain-level associations within individual studies (Supplementary Tables 4-7), including several strains belonging to the *Bacteroides* genus within the rare cancer cohort, which has been previously implicated in responses to CTLA-4 blockade(45).

These findings reinforce the conclusion that the lack of reproducible microbiome taxonomic markers across studies is unlikely to be attributable to analytical choices(28), as StrainSpy identified both well-established and novel associations within the CRC dataset. However, these approaches do not account for gene-level variation and may therefore miss associations driven by functional differences in the microbiome that are less dependent on individual species or strains, such as those underlying short-chain fatty acid production(46, 47). Recent studies have highlighted the diet-gut axis as a major determinant of ICI response. For example, obesogenic diets have been shown to restore ICI sensitivity following a shortterm dietary switch(48). Given the well-established influence of diet on gut microbiome composition(49, 50), differences in dietary patterns across study populations could also potentially account for a substantial proportion of the heterogeneity observed between studies.

Taken together, StrainSpy allows for the integration of large multi-cohort microbiome datasets into a single model, accounting for compositional and batch effects while identifying strain-level associations missed by conventional abundance-based methods. The recovery of known and novel associations in CRC, alongside the lack of taxonomic signals across immunotherapy cohorts, consistent with previous meta-analyses (28, 32), further supports StrainSpy’s low false-positive rate and ability to distinguish true biological signals in noisy datasets.

### Containment ANI features identified by StrainSpy achieve predictive performance as accurate as abundance-based approaches

Given the interest in microbial features for machine learning-based diagnostics, we next tested whether StrainSpy-identified cANI features could allow prediction of complex host phenotypes, such as disease status and response to therapy. As cANI captures strain-level genomic variation while being robust to compositional effects, these features may provide paths to more stable biomarkers with potential for translation into targeted assays such as qPCR.

Similar to the previous section, we evaluated the viability of this approach using both the large CRC collection and the immunotherapy microbiome cohorts. To assess model performance, we used leave-one-dataset-out cross-validation, whereby each dataset was iteratively held out as an independent test set. For each training iteration, we first identified significant strain-level features using StrainSpy and subsequently trained an elastic net machine learning classifier using these features.

As seen in Figure 5a, the cANI-based models demonstrated strong predictive performance in leave-one-dataset-out (LODO) analysis, with AUCs ranging from 0.70 to 0.93. This closely mirrors the 0.71 to 0.95 range reported in previous abundance-based efforts by Piccinno et al(24). Only significantly associated strains identified within each training fold were used as features for model training (see Methods). To further evaluate model generalisability, we performed cross-cohort prediction analyses, training models on individual cohorts and evaluating performance on independent cohorts (see methods, Supplementary Figure S5). Similar to the previous analysis, the average and best AUC values for each cohort were within the range of the original meta-analysis(24), illustrating that strain-level predictors retained predictive capacity across studies. Although direct comparison is limited by differences in training cohorts and classifiers (publicly available samples and elastic net in our study versus Random Forest in the original study), the similar performance demonstrates that cANI-derived features capture reproducible CRC-associated signals across heterogeneous cohorts and can achieve predictive accuracy comparable to conventional abundance-based methods.

**Fig. 5.**
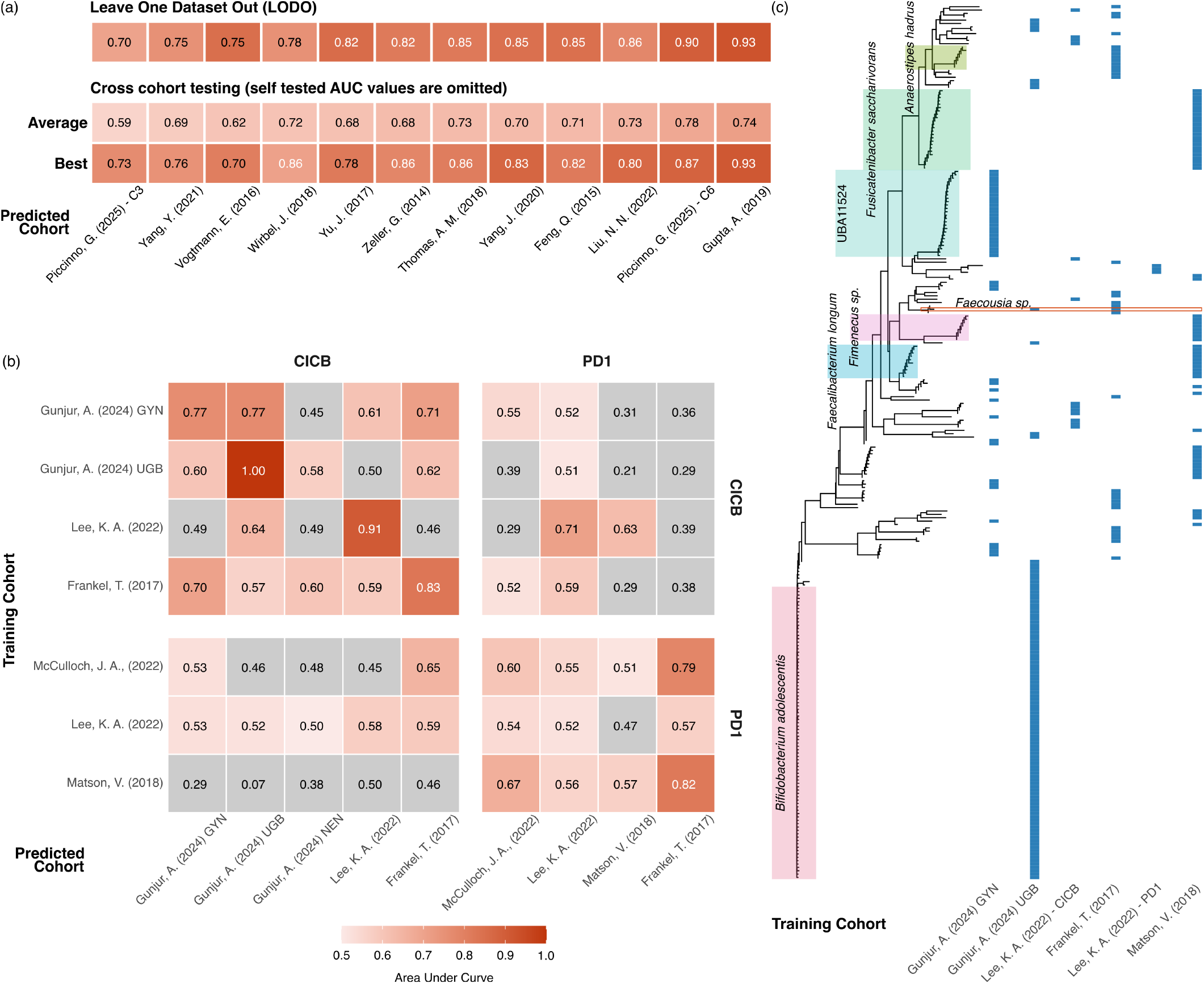
ANI predictions achieve similar prediction performance to abundance based predictors. (a) Summary of prediction performance across the pooled colorectal cancer cohorts. The first row shows AUC values from the LODO analysis, second and third rows show the average and best AUC values in cross-cohort prediction analysis. (b) Cross-cohort prediction performance across the immunotherapy datasets. The Melanoma cohorts were stratified by treatment regimen where applicable. Only models with >20 samples were used as a training cohort. (c) Phylogenetic distribution of strain-level features selected independently across immunotherapy cohorts. Some species with a large number of strains are highlighted on the tree. Blue tiles indicate whether the strain was selected in the corresponding training cohort. Only a single strain (highlighted in red) was independently identified in more than one cohort, illustrating the limited overlap in strain-level microbiome markers across cohorts.

Next, although we did not identify cross-cohort associations within the immunotherapy dataset, we tested whether strains associated with response in one cohort could predict immunotherapy sensitivity in an independent cohort. Given the influence of treatment modality (CTLA-4 blockade versus PD-1 inhibition) on outcomes, melanoma co-horts were stratified by treatment type where applicable. Within the pancancer dataset, only the GYN and UGB cohorts had sufficient sample sizes (>20) for model training. As seen in Figure 5b, an elastic net model trained on GYN-associated strains achieved AUCs of 0.77 in UGB and 0.45 in NEN, while a model trained on UGB-associated strains achieved AUCs of 0.6 in GYN and 0.58 in NEN. Performance across melanoma cohorts, using features derived from both pancancer and melanoma datasets, was variable (AUC 0.21–0.82), consistent with the cohort-specific heterogeneity observed in the association analyses (Figure 5c). These results are generally consistent or slightly improved from previous abundance-based analyses(32). Overall, we found that StrainSpy effectively selects cANI features for downstream machine learning models, achieving comparable accuracy to abundance-based approaches.

## Discussion

Strain-level genetic variation represents an important but underutilised source of microbiome variation, as closely related strains can exhibit distinct functional capacities and associations with host phenotypes. StrainSpy is a flexible statistical algorithm for identifying strain-level associations across a wide range of microbiome study designs using cANI. By combining cANI with a zero-inflated Beta model and empirical Bayes priors, StrainSpy accounts for both variation in strain presence-absence and differences in cANI among detected strains. This enables robust inference across cohorts of varying sizes while mitigating challenges associated with compositional microbiome data.

Through extensive simulations and analyses of well-characterised metagenomic datasets, we demonstrate that StrainSpy consistently identifies intra-species genetic varia-tion with accuracy comparable to, or exceeding, that of existing strain-identity-based approaches. In particular, our *Escherichia* sp. spike-in simulation illustrates a practical distinction between cANI-based and phylogeny-based approaches like AnPan. While phylogeny-based methods can provide highly resolved strain inference when sufficient sequence coverage is available, their application depends on successful marker extraction and reconstruction of a reliable strain phylogeny. In contrast, cANI-based approaches can operate directly on genome similarity estimates without requiring a species-specific phylogeny.

Importantly, a re-analysis of a longitudinal antibiotic perturbation study demonstrated the advantages of using cANI rather than conventional species-level abundance analyses. Strain-level analyses revealed substantial within-species perturbations, including in *V. hominis* and *B. uniformis*, that would have been obscured by species-level approaches. Previously reported reductions in *E. coli* abundance were found to be potentially confounded by the overall decrease in microbial load following antibiotic administration, whereas StrainSpy was robust to this effect.

A major strength of StrainSpy is the use of a generalized linear mixed model framework. Coupled with cANI, StrainSpy enables straightforward integration of large, multi-cohort microbiome studies. Re-analysis of large cancer microbiome meta-studies demonstrated that StrainSpy recovered previously established associations, such as *Fusobacterium animalis* (*Fusobacterium nucleatum* subsp. *animalis*) in colorectal cancer, while also identifying novel strain-level associations, including within *E. coli*, that were not detected by alternative approaches. Importantly, StrainSpy identified no significant strain level associations in a meta-analysis of microbiome correlates of immunotherapy response. This is consistent with the low false-positive rate observed in our simulations and congruent with previous reports suggesting limited reproducibility of microbiome taxonomic predictors across immunotherapy cohorts(28). A possible explanation is that functional differences in the microbiome which interact with immunotherapy, such as short-chain fatty acid production, may be less dependent on individual species or strains.

Beyond association testing, we also demonstrated that StrainSpy can identify cANI-based features for use in downstream machine learning models. In both the colorectal cancer and immunotherapy cohorts, models built using StrainSpy-selected features achieved predictive performance comparable to conventional abundance-based approaches while requiring fewer features. Because detecting the presence or absence of specific strains or genetic markers is often simpler than performing whole-community sequencing to accurately quantify species abundances, these features could facilitate the development of simpler and more cost-effective microbiome based diagnostic assays, such as qPCR-based tests.

While StrainSpy avoids many of the challenges associated with modelling abundance data, it cannot detect microbiome functional dynamics that occur without a change in the ANI of a strain or species. Consequently, we view StrainSpy and other similar methods, as complementary to, rather than a re-placement for, existing abundance-based methods. Although StrainSpy circumvents certain batch effects, such as those introduced by changes in microbial load, identity-based batch effects such as strain compositional differences between populations remain a challenge. For instance, distinct strains or lineages circulating in different geographical regions can still introduce bias. It is therefore crucial to include potential batch effects within StrainSpy’s GLMM framework. Finally, while StrainSpy identifies associations between ANI and covariates of interest, it does not reconstruct strain haplotypes or explicitly model the underlying population structure of a species. Here, phylogenetic based approaches, such as An-Pan, can provide complementary insights into the populations underlying these associations.

Overall, StrainSpy provides a powerful framework for characterising strain-level variation in microbiome studies, complementing existing abundance-based analyses and enabling the detection of associations that may be obscured by conventional species-level analyses.

## Methods

### Strain and phenotype data

The primary input to StrainSpy is a SummarizedExperiment object(51) containing both strain and phenotype data. Generally, strain data is a table of containment average nucleotide identity (ANI) estimates between a set of metagenomic samples and a reference collection of genomes. In practice, the references can be generated by deduplicating Genome Taxonomy Database (GTDB)(52) at approximately 99% nucleotide identity in order to capture strain-level diversity. Alternatively, a broader species-level set of genomes can be extracted by deduplicating at 95% identity. Usually, containment ANI estimates can be obtained using Sylph(17) in query mode. Alternatively, Sylph profile mode or outputs from other profiling tools such as MetaPhlAn(53) or Sourmash(18) can also be used. Strains that are largely absent across samples should be removed prior to analysis. For cross-sectional data, StrainSpy has a default filter to exclude strains that are observed in <10% of samples. For longitudinal studies, filtering should consider repeated observations within the same subject, retaining strains that appear in at least a minimum number of subjects at multiple time points.

### Model fitting

The general model in StrainSpy has the form: ANI ~ Phenotype + Covariates. This approach differs from previous cANI based methods, which model disease status (or another phenotype) as the outcome and the microbial features (relative abundance or ANI) as covariates. Modelling microbial features as outcomes permits likelihoods that directly reflect the biological mechanisms underlying strain turnover. For cANI, disease-associated changes may manifest as complete disappearance of a strain, replacement by a closely related strain, or a combination of both. Furthermore, because containment ANI is a bounded quantity with excess zeros and a continuous variation near identity, modelling it as the outcome enables direct representation of strain depletion (inflation of zeros) and replacement (shifts in ANI). In a typical analysis, StrainSpy fits several thousand univariate models, rendering a fully Bayesian approach computationally infeasible for most studies. Instead of maximum likelihood estimation, StrainSpy uses maximum a posteriori (MAP) estimation with weakly informative priors as the default approach as implemented in the glmmTMB R package(54, 55). Complete details on model fitting and available model options are provided in Supplementary Methods.

#### Zero inflated beta model

While Strainspy does not impose a restriction on the choice of model family, the ZiB model is used as the default as it accounts for both the inflated number of zeros driven by the absence of strains and the distribution of cANI which is bounded by 1 indicating an identical strain (within the detection limit) to the reference(56). The ZiB model decomposes the data-generating process into two components: a logistic submodel governing the probability of observing a zero value, and a beta-regression submodel which accounts for non-zero ANI values on the interval (0.95-1). In practice, cANI values are offset by 0.01 to mitigate edge effects near the upper boundary.

By jointly modelling depletion and identity variation within a single likelihood, ZiB aggregates evidence across both components when testing for associations. This can improve statistical power, particularly when disease effects are distributed across partial strain depletion and replacement.

#### Ordered beta regression model

The ordered beta regression(36) model provides an alternative likelihood for bounded continuous responses that naturally accommodates observations at the boundaries of the unit interval (i.e. cANI range 0.95 - 1). Unlike the ZiB model, it does not explicitly decompose the response into presence/absence and continuous components. Instead, all observations are modelled within a single likelihood, making the model appropriate when strains are consistently detected across samples and the biological signal is driven primarily by changes in strain identity rather than strain depletion.

In practice, ordered beta regression is recommended for datasets where the majority of strains are present in most samples, such as longitudinal studies or analyses restricted to consistently detected taxa. Under these conditions it can provide greater statistical power than ZiB because model complexity is reduced and inference focuses directly on continuous variation in cANI.

#### Logistic regression model

StrainSpy also implements the logistic regression framework proposed by Shaw and Yu (17) to facilitate comparison with previous cANI-based methods. In this formulation, phenotype is treated as the response and cANI as the predictor after thresholding cANI values to define strain presence. Unlike the Generalised-Linear Mixed Modelling (GLMM) framework used throughout StrainSpy, this approach cannot simultaneously model strain depletion and replacement because continuous cANI values below the chosen threshold are discarded. Consequently, inference depends on a user-defined threshold that determines strain presence. Simulation studies demonstrated that sensitivity varied substantially with threshold choice, indicating that logistic regression may fail to detect biologically relevant strain variation depending on the chosen threshold (see Supplementary Methods for details).

#### Empirical Bayes priors

Empirical Bayes priors are estimated by borrowing information across all strain-specific models. Rather than specifying identical priors for every strain, StrainSpy first fits simplified models independently for each strain and uses the resulting coefficient distributions to estimate population-level prior distributions.

The advantage of using a MAP estimation framework(55) is the ability to shrink the strain-wise variance estimates towards the global variance, reducing unstable estimates and limiting the number of false positives.

Separate prior distributions are estimated for the zero-inflation and beta components because these represent distinct biological processes. For the zero-inflation component, cANI values are first binarised into detected and undetected strains, and logistic regression(57) is applied independently for each strain. The resulting coefficients are pooled across strains to estimate the centre and spread of the prior distribution.

Similarly, a fast regression approach(58) is used on the beta component using only samples in which the strain is detected. The resulting empirical distributions are used as Gaussian priors during MAP estimation, shrinking unstable coefficient estimates towards values supported across the full dataset.

### Incorporating taxonomy

Taxonomic data for each strain in the database can be provided as input to strainspy for various downstream analyses, including taxonomy-aware multiple testing correction, grouping of associated strains, and visualisation. Taxonomy provides a hierarchical framework for organising strain-level associations and enables results to be interpreted in the context of related microbial lineages.

For the analyses presented in this manuscript, taxonomic annotations were obtained from GTDB(52) and provided alongside the reference genome catalogue through Zenodo (see Data Availability). For user-defined genome databases, taxonomic information can be supplied as a tab-separated file containing one row per reference strain and columns corresponding to the relevant taxonomic ranks.

When a curated reference database contains the exact strain of interest for a population, taxonomic annotations can provide a direct biological interpretation of detected associations. However, for broad genome catalogues such as GTDB, associated reference genomes should be interpreted in the context of the available reference diversity, as the exact strain present in a sample is unlikely to be represented in the database.

#### Correction for multiple testing

By default, StrainSpy performs multiple testing corrections at the strain level using the Benjamini–Hochberg method, assuming each strain is independent. Alternative corrections (Bonferroni, Holm, Hochberg, and Hommel) are also available.

StrainSpy can also incorporate the taxonomy of strains present in a database by using the harmonic p-value method(33) to aggregate strains at higher taxonomic levels (e.g., species or genus). This allows multiple partially similar strains to collectively contribute to a higher-level signal, thereby revealing associations that may be missed when considering strains individually.

### Simulations

### Using signal implantation to simulate strain identity variation and depletion

We adopted a signal implantation strategy to introduce controlled strain-level effects directly into metagenomic data, rather than relying on parametric simulators. This approach has been shown to better reproduce metagenomic data characteristics while maintaining realistic disease effect sizes(16). A random subset of 200 metagenomes from Zeevi *et al*.(35) (accessions in Supplementary table 8) was split equally into cases and controls. Strain profiling was done using Sylph v.0.9.0(17) in query mode against a custom GTDB database dereplicated at 99% identity (see Data Availability). After removing strains observed in <10% of metagenomes, 18,027 strains with ANI > 95% remained, from which 100 were randomly selected for modification in cases. Identity variation was simulated by scaling ANI by a factor: SF ∈ [0.975, 1], corresponding to an ANI reduction up to 2.5%, while depletion was simulated by setting ANI = 0 in a randomly selected subset of up to 75% of non-zero samples.

#### Spiking reads from Escherichia sp. GCF_009820115.1 and Parabacteroides distasonis GCF_024791025.1

Baseline metagenomes were first profiled using Sylph v.0.9.0(17) to characterize the strain composition of the simulation cohort (see Supplementary Table 8). For each target species, pairwise ANI(59) comparisons were calculated between genomes in the GTDB reference database to characterize the genetic relationships among candidate strains. Candidate spike-in genomes were selected from strains that were not represented among the dominant strain-level signals in the baseline metagenomes, or were sufficiently divergent from strains detected in the baseline cohort. This ensured that the introduced genomes represented novel strain-level variation relative to the existing population structure rather than increasing the abundance of an already observed strain.

Following strain selection, samples for spike-in were chosen based on the observed distribution of the corresponding species in the baseline cohort. For the *Escherichia* sp. spike-in experiment, 20 metagenomes were randomly selected from the cohort for introduction of the simulated strain. For the *P. distasonis* spike-in experiment, since our motivation was to compare StrainSpy with AnPan(21), sample selection was restricted to the subset of samples in which *P. distasonis* was detected by StrainPhlAn(39). Specifically,*P. distasonis* was only identified in 100 of the 200 baseline metagenomes, and 20 samples were randomly selected from this subset for spike-in experiments. This design enabled evaluation of strain-level association methods under condi-tions where the introduced genome represented additional strain diversity within an existing species population.

We used art_modern v.1.3.4(37), based on art 2.5.8(38) to simulate the reads from each genome. Paired end reads were simulated at a fold coverage of 10*×*. For each spike-in experiment, a random portion of these reads were randomly subsampled to achieve the desired effective coverage. The *Escherichia* sp. spike-in was performed at a fixed coverage of 1x, where 10% of reads were randomly sampled. For *P. distasonis*, low (1*×*), medium (3*×*) and high (5*×*) coverage scenarios were achieved by sampling 10%, 30% and 50% of reads, respectively. Samples chosen as controls and cases for the two spike-in simulations are available in Supplementary Table 8.

### Testing for associations in gut metagenomic datasets

For each dataset, strain-level associations were evaluated using dataset-specific fixed and random effects reflecting the original study design. In the longitudinal antibiotic perturbation dataset(25), time following antibiotic treatment (days) was modelled as a discrete fixed effect with subject included as a random intercept to account for repeated sampling: ~days + (1 | subject). An alternative model additionally adjusted for microbial load was also fitted: ~days + (1 | subject) + load. In the pooled colorectal cancer dataset(24), disease status was modelled while adjusting for age, sex and body mass index (BMI), with study included as a random intercept to account for between-cohort heterogeneity. For the immunotherapy cohort(32), response to treatment (responder versus non-responder) was modelled while adjusting for histology, age, sex and BMI. For melanoma cohorts (26– 31), response to treatment (responder versus non-responder) was modelled with and without adjusting for type of therapy (CICB versus PD-1). All metadata and sample accessions for these datasets are provided in Supplementary Tables 9-12.

### Searching for *E. coli* adhesin genes in CRC metagenomes

Reference sequences of *E. coli* adhesin genes were extracted from the adhesiomeR package(60). ARIBA v.2.14.7 (61) was used to identify these reference sequences directly from the metagenomic reads. Only paired-end metagenomes from control and CRC samples in which *E. coli* was detected by Sylph(17) were used in the analysis. This allowed us to examine differences in adhesin carriage among *E. coli*-positive samples without being biased by the variation in overall carriage of *E. coli*. For each adhesin gene, we compared its prevalence between CRC and control groups using a binomial generalized linear mixed model(62, 63), with case status as the response, gene presence/absence as the explanatory variable, and study included as a random effect to account for between-study variation. Genes found in <25 samples (approximately 10% of the population) were excluded from analysis and p-values were adjusted for multiple testing using the Benjamini-Hochberg procedure.

### Phenotype Prediction

All training and prediction tasks were performed using the R package caret v.7.0.1 (64) using the elastic-net regression model implemented in the R package glmnet v.5.0 (65).

### Pooled colorectal cancer dataset

For the pooled colorectal cancer dataset(24), leave-one-dataset-out (LODO) analysis was performed by iteratively withholding each dataset as an independent test set. In each iteration, StrainSpy ZiB models were fitted after filtering strains with less than 10% nonzero cANI values. For all training datasets the fitted model had the form ~cANI disease + age + sex + BMI + (1 | study), with empirical Bayes priors. Strain-level features significant at an adjusted p ≤ 0.05 using the Benjamini-Hochberg procedure were selected from the training data and used as predictors for elastic-net model training (see Supplementary table 13). Predictive performance was evaluated on the held-out dataset using the area under the receiver operating characteristic curve (AUC).

The portability of strain-level predictors across independent cohorts was assessed using a cross-study prediction analysis. Each dataset was independently used as a training cohort. After applying the 10% prevalence filter, StrainSpy ZiB models of the form ~cANI disease + age + sex + BMI were fitted separately within each training cohort using empirical Bayes priors. Because individual datasets had substantially smaller sample sizes compared to pooled cohorts, this resulted in reduced power for multiple-testing corrected feature discovery. Therefore, strain features were ranked by association strength rather than filtered using a significance threshold. The number of predictors was fixed to match the pooled analysis (6655), and the top-ranked strains were used for elastic-net model training (see Supplementary table 14). Models were then evaluated by predicting CRC status in each independent dataset using AUC.

#### Response to immunotherapy in pancancer

Cross-cohort prediction of immunotherapy response was performed using the same elastic-net framework described above. Cohorts were analysed independently, with samples separated by cancer type for rare tumour cohorts and by dataset and treatment modality (CICB versus PD-1 therapy) for melanoma cohorts. Within each cohort, strain-level associations with treatment response were evaluated using StrainSpy ZiB models after applying the 10% prevalence filter.

Strain-level features associated with response within each training cohort were selected using Benjamini–Hochberg adjusted significance thresholds and used as predictors for elastic-net model training. Supplementary Table 15 details the number of strains used for each model. For the cohorts that did not yield significant strain-level associations after multiple-testing correction, strain features were ranked by association strength based on p-value, and the top 100 ranked strains were selected as predictors. Models were subsequently evaluated by predicting responder versus nonresponder status in independent cohorts using AUC.

#### Estimating the microbial load

We used Microbial Load Predictor (MLP)(14) with the Galaxy model to predict the microbial load in all samples collected in Palleja *et al*.(25). Predicted load was log transformed before using it as a covariate in association testing.

## Data and code availability

The genome dereplication workflow used to generate the custom databases is available at https://github.com/gtonkinhill/genoreps. The resulting 99% ANI Sylph database used in this study has been deposited in Zenodo (DOI: 10.5281/zenodo.21796878), available https://zenodo.org/records/21796878. All genome accessions and metadata used for downstream analyses are provided in Supplementary Tables 8-12, and the complete analysis code is available at https://sudaraka88.github.io/strainspy-manuscript/.

## Acknowledgements

This work was supported by resources provided by the Pawsey Supercomputing Research Centre’s Setonix Supercomputer (https://doi.org/10.48569/18sb-8s43) and Acacia Object Storage (https://doi.org/10.48569/nfe9-a426),with funding from the Australian Government and the Government of Western Australia. Funding was provided by the Australian Research Council (grant no. DE240100316 to G.T.-H. and DE220100965 to V.R.M.), the National Health and Medical Research Council (grant no. GNT2025515 to G.T.-H., GNT2041653 to D.J.I. and 2036149 to S.S., V.R.M,S.B. and K.T.) and the Spanish Ministry of Science and Innovation (RYC2023-042907-I to V.R.M.).

## Ethics, Consent to Participate, and Consent to Publish declarations

Not applicable.

## Supplementary Methods

### Model fitting

StrainSpy performs association testing by fitting a collection of strain-specific generalized linear mixed models (GLMM), allowing flexible specification of both response distributions and study-specific fixed and random effects.

Let i = 1, …, n denote samples and j = 1, …, p denote strains. The response matrix **Y** = (y_ij_) contains the containment average nucleotide identity (cANI) between sample i and strain j. cANI is defined on the bounded interval [0, 1], where y_ij_ = 0 indicates that strain j was not detected in sample i, and non-zero values typically lie in the interval [0.95, 1]. Although y_ij_ = 0 generally reflects biological absence, it may also arise from limited sequencing depth or imperfect strain assignment.

For each strain j, the response vector **y**_j_ is modelled as

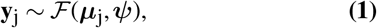

where *F* denotes the chosen response distribution, ***µ***_j_ is the conditional mean, and ***ψ*** represents any additional distribution-specific parameters such as dispersion or zero-inflation parameters. The conditional mean: ***µ***_j_ = E (**y**_j_ | **b**_j_) is related to the linear predictor:

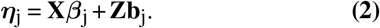

via the invertible link function g(***µ***_j_) = ***η***_j_.

Here, **X** is the fixed-effect design matrix containing pheno-type and optional sample-level covariates, and ***β***_j_ is the corresponding vector of strain-specific fixed-effect coefficients. When specified, **Z** is the random-effects design matrix, and **b**_j_ denotes the strain-specific random effects.

StrainSpy treats strain identity (cANI in this case) as the re-sponse variable. This concept follows the hypothesis that the host phenotype such as case control status, or external perturbations such as drug treatment or dietary intervention, alter microbial strain composition, and naturally leads to both strain depletion and strain replacement within a unified probabilistic framework.

### Parameter estimation

Since StrainSpy fits a large number of independent strain-specific GLMMs, fully Bayesian inference is computationally challenging due to the need to repeatedly integrate over high-dimensional latent structures. Furthermore, unregularised maximum likelihood or restricted maximum likelihood estimation can lead to unstable parameter estimates in sparse data settings, particularly in the presence of separation and strong correlations among predictors (1), which are common characteristics of strain-level metagenomic data.

#### MAP estimation

To address this, StrainSpy uses maximum a posteriori (MAP) estimation as implemented in the R package `glmmTMB` (2).

Briefly, for each strain j, let *θ*_j_ = (***β***_j_, **b**_j_) denote the full set of fixed and random effect parameters. The model is estimated by maximising the penalised log-likelihood:

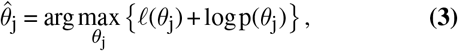

where *l* (*θ*_j_) denotes the GLMM log-likelihood and p(*θ*_j_) represents prior distributions that induce regularisation on model parameters.

This regularisation improves numerical stability in highdimensional and sparse settings, and mitigates issues such as complete or quasi-complete separation and unstable variance component estimation that commonly arise in strain-level association studies.

In practice, weakly informative priors are used by default to balance stability and sensitivity in association testing. Alternatively, stronger priors are available for stricter control of false positives. Optionally, an empirical Bayes method is provided for tighter control, especially when analysing smaller datasets.

#### Empirical Bayes estimation of MAP priors

Instead of relying solely on fixed weakly informative priors, StrainSpy can optionally estimate prior distributions empirically from the data.

First, for the zero-inflation component, cANI values are transformed into a binary indicator of strain presence:

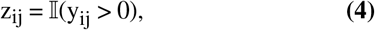

where z_ij_ = 1 indicates that strain j is detected in sample i, and z_ij_ = 0 indicates non-detection.

A binomial GLM is then fitted to model the probability of strain detection:

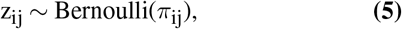

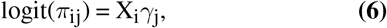

where X_i_ is the vector of fixed-effect covariates for sample i (e.g. phenotype and other sample-level covariates), *γ*_j_ is the corresponding strain-specific coefficient vector, and *π*_ij_ denotes the probability that strain j is present in sample i. Next, for the non-zero component, for each strain, only observations with y_ij_ > 0 are retained, corresponding to samples in which strain j is detected. The conditional distribution of cANI values is modelled using a beta regression:

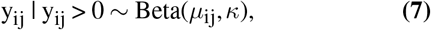

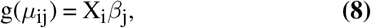

where *µ*_ij_ is the conditional mean of the beta distribution, *κ* is a dispersion parameter controlling variability around the mean, and *β*_j_ is the strain-specific coefficient vector associated with the fixed-effect design matrix X_i_.

Parameter estimates from both components are then aggre-gated across strains to estimate empirical distributions of the corresponding coefficients. These empirical distributions are used to define updated prior specifications, inducing adaptive shrinkage of strain-specific coefficients towards a shared population-level distribution.

This shrinkage reduces variance in parameter estimates while preserving strain-specific signals, improving robustness in sparse, high-dimensional settings typical of metagenomic strain profiling, particularly in datasets with limited sample sizes (around 30).

### Likelihood families

StrainSpy supports multiple likelihood families specified in `glmmTMB`.

#### Zero-inflated beta model

The zero-inflated beta (ZiB) model is set as the default likelihood since we have found it is well suited to model the distribution of cANI. To ensure the numerical stability of the beta component of the model, cANI values are bounded away from the limits of the unit interval. Specifically, observed values are transformed as:

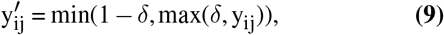

with *δ* = 0.01. This prevents boundary issues in the beta likelihood and stabilises parameter estimation. This likelihood is modelled as:

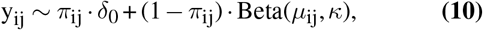

where *δ*_0_ denotes a point mass at zero, *π*_ij_ is the zero-inflation probability, and *κ* is a dispersion parameter.

The two components are modelled as:

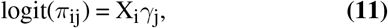

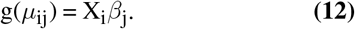

This formulation jointly captures strain depletion (zeroinflation component) and strain replacement (beta component).

#### Ordered beta regression model

The ordered beta regression model is used for modelling continuous outcomes bounded in the interval [0, 1] and is particularly suited to settings where zero-inflation is not present but strong boundary effects may still occur. Unlike standard beta regression, the ordered beta formulation is derived from a latent variable representation that maps a continuous response through a flexible ordered structure, avoiding numerical and interpretability issues associated with boundary behaviour (3).

In StrainSpy, we have found this model can be effective when users are interested in detecting changes in strain identity (cANI variation) among strains that are consistently present across samples, rather than changes in strain prevalence.

The ordered beta model provides a more flexible likelihood than standard beta regression by relaxing strict distributional assumptions near the boundaries, leading to improved robustness when modelling high-identity genomic similarity measures such as cANI. For a more complete description of this model see Kubinec et al., 2023 (3).

### Logistic regression approach

Following the MWAS framework described in (4), StrainSpy includes an association model in which strain-level cANI is used as a predictor of a binary phenotype.

Let i = 1, …, n denote samples and j = 1, …, p denote strains. The binary phenotype z_i_ ∈ {0, 1} (e.g. case/control status) is modelled using logistic regression:

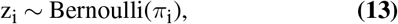

with linear predictor:

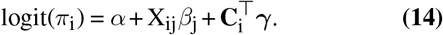

Here, the model is fit independently for each strain j, where X_ij_ denotes the cANI value for sample i and strain j, and **C**_i_ represents additional sample-level covariates such as age and sex.

Following the original approach, high-identity strain variation is inferred by applying a thresholded transformation on cANI:

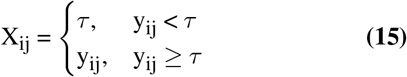

where *τ* is a fixed threshold (e.g. *τ* = 0.98 used in (4)). This shrinks all low cANI values to a constant while preserving variation among high-identity matches.

A key motivation for this approach is to omit less informative cANI values from association testing. However, as demonstrated in simulation studies, while certain thresholds can yield highly accurate performance, the resulting inference is sensitive to the choice of *τ*, and has an impact on both sensitivity and specificity (see Supplementary Figure S6).

## Supplementary Figures

**Fig. S1.**
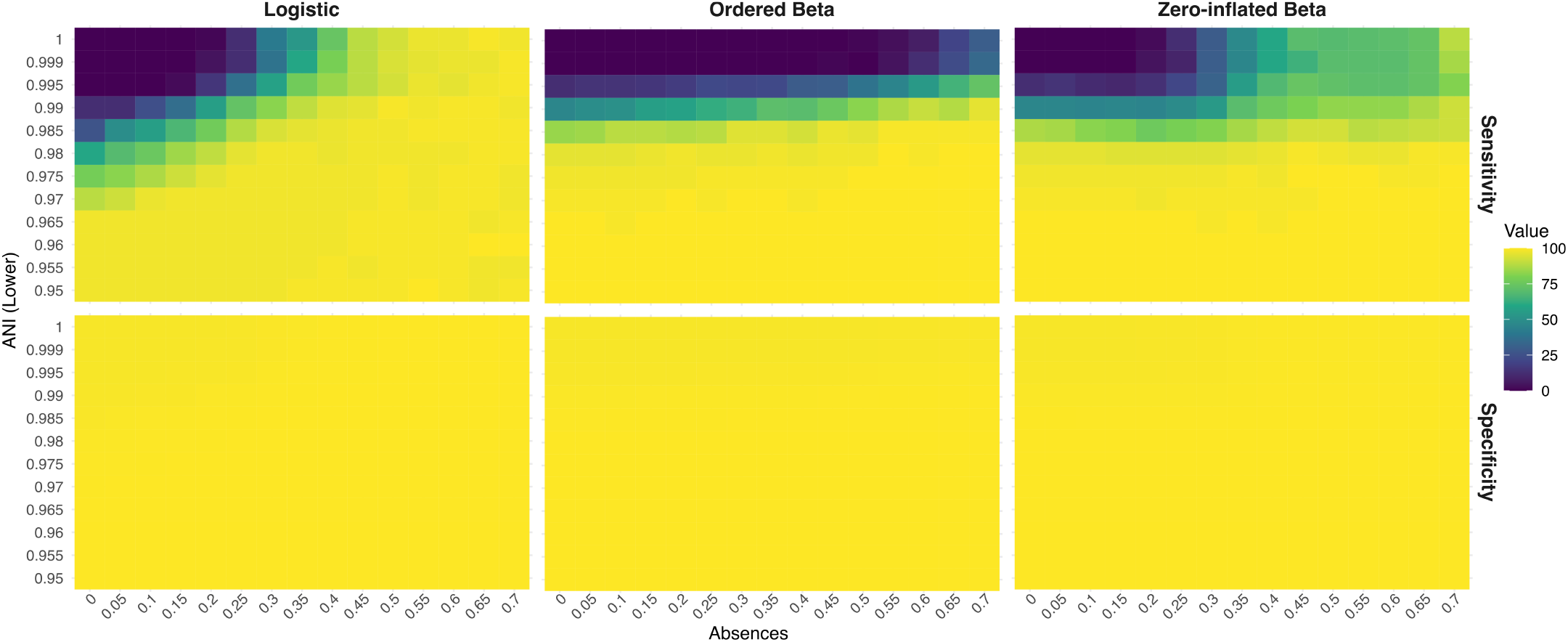
Performance of StrainSpy in simulated case-control simulations. Heatmaps show the sensitivity (top) and specificity (bottom) of logistic regression (left), ordered beta (middle), and zero-inflated beta (ZiB; right) models across a range of strain depletions (x-axis) and reductions in average nucleotide identity (ANI; y-axis). Each heatmap cell corresponds to a unique combination in which 100 randomly selected strains were depleted (i.e., ANI = 0) in the indicated proportion of 100 randomly selected case samples (by the proportion shown in the x-axis) and/or had their ANI reduced (by the indicated amount y-axis), compared with 100 control samples. This extends the simulations shown in Figure 3a by evaluating combinations of identity variation and strain absence simultaneously. Overall, ZiB maintained the highest sensitivity across most mixed scenarios while preserving high specificity, whereas logistic regression was generally less sensitive and the ordered beta model performed poorly when strain depletion was substantial. For all logistic regression models, the ANI threshold was set to τ = 0.97 (see Supplementary Methods).

**Fig. S2.**
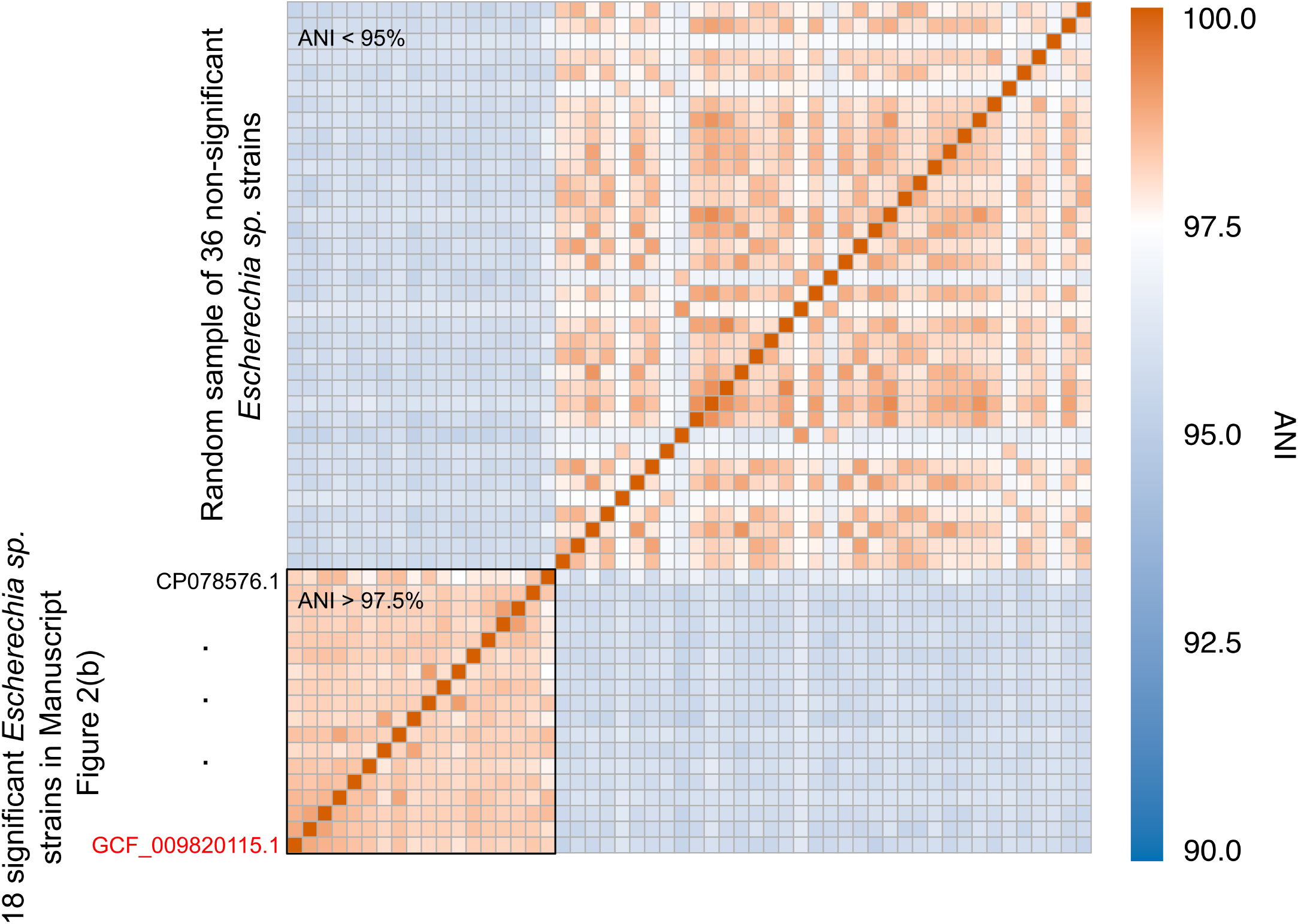
Pairwise ANI between significantly associated and randomly selected non-significant *Escherichia* sp. strains in the spike-in simulation. Heatmap showing pairwise ANI, measured using skani (see legend), between the 18 significantly associated strains and 36 randomly selected non-significant *Escherichia* sp. strains, ordered from bottom to top and left to right. The 18 significant strains are ranked by significance as shown in Figure 3b in the manuscript. These strains form a highly similar cluster with pairwise ANI > 97.5%, explaining why simulated reads from GCF_009820115.1 increased cANI for all 18 strains. In contrast, the randomly selected *Escherichia* sp. strains exhibit substantially lower ANI with the significant cluster and show a heterogeneous distribution of pairwise ANI values among themselves.

**Fig. S3.**
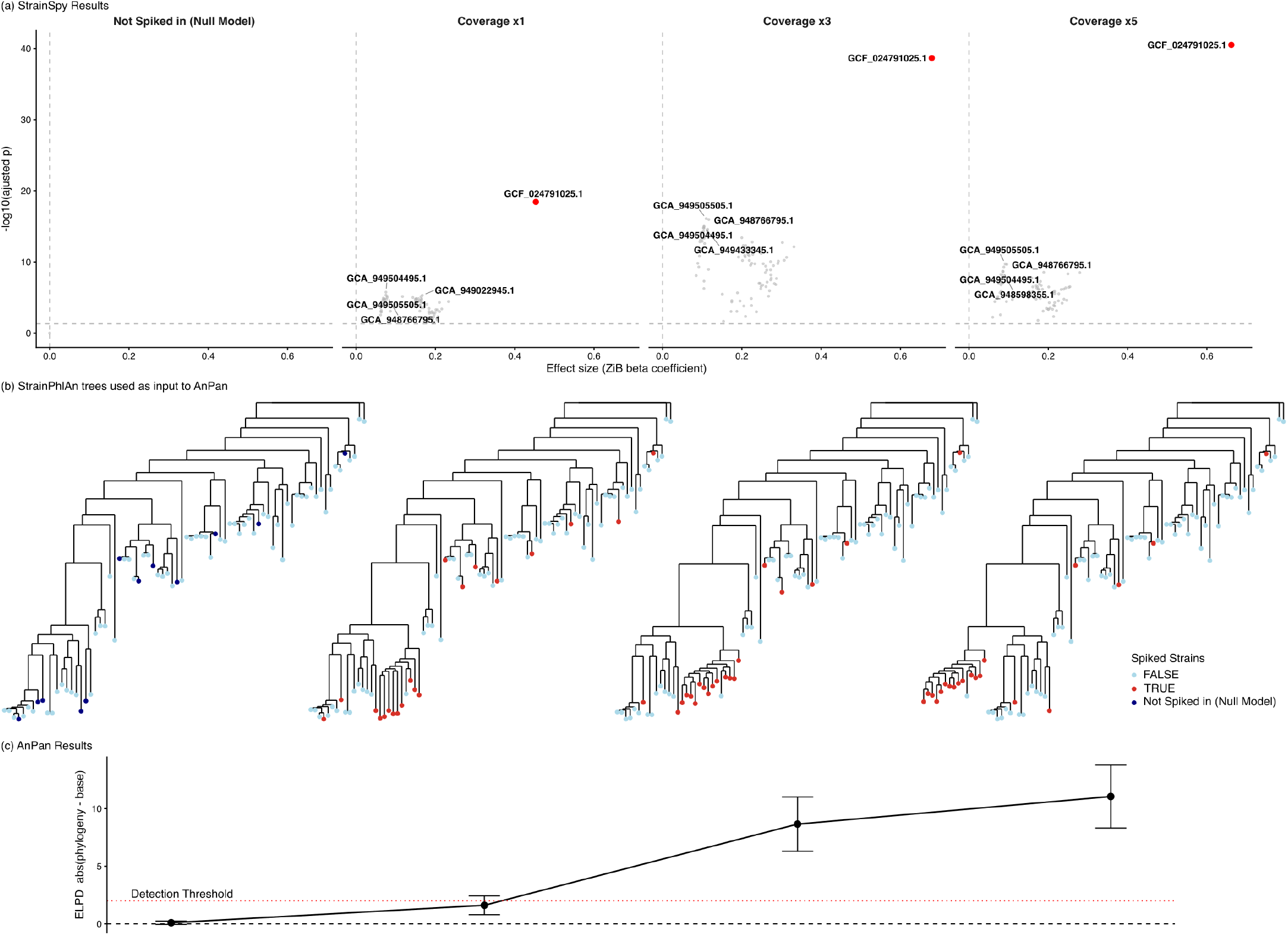
Comparison of StrainSpy and Anpan under simulated spike-in conditions at varying coverage levels. (a) StrainSpy differential association results (ZiB model) for simulations with no spike-in (null), and spike-in at 1*×*, 3*×*, and 5*×* coverage in 20 randomly selected *Parabacteroides distasonis* (SGB1934) samples. Volcano plots show model significance across strains, with the spiked strain (GCF_024791025.1) consistently identified as the top hit in all non-null simulations (adjusted p-values of 3.68 *×* 10^−19^,2.21 *×* 10^−39^, and 3.11 *×* 10^−41^, respectively). Additional closely related *P. distasonis* strains are also detected at lower significance due to high ANI, similar to the *Escherichia* sp. spike-in simulation. As expected, no significant associations are observed under the null, where no spike-in reads are present. (b) StrainPhlAn phylogenetic trees used in the Anpan analysis. Tips coloured by spike-in status. Red - true spiked-in, and blue: non-spiked strains. In the null, no true spike-in signal is present; however, a similar number of strains are randomly designated as “spike-in” to match the structure required by the comparative model, serving as a negative control. At 1*×* coverage, Anpan did not detect a phylogenetic signal. However, at 3*×* and 5*×* coverage, the spiked strain is reliably recovered within the inferred clade structure. (c) Anpan model comparison showing the difference in expected log predictive density between phylogenetic and non-phylogenetic models. At 1*×* coverage, the difference is below the detection threshold. At higher coverage, the phylogenetic model is increasingly favoured, with larger positive differences.

**Fig. S4.**
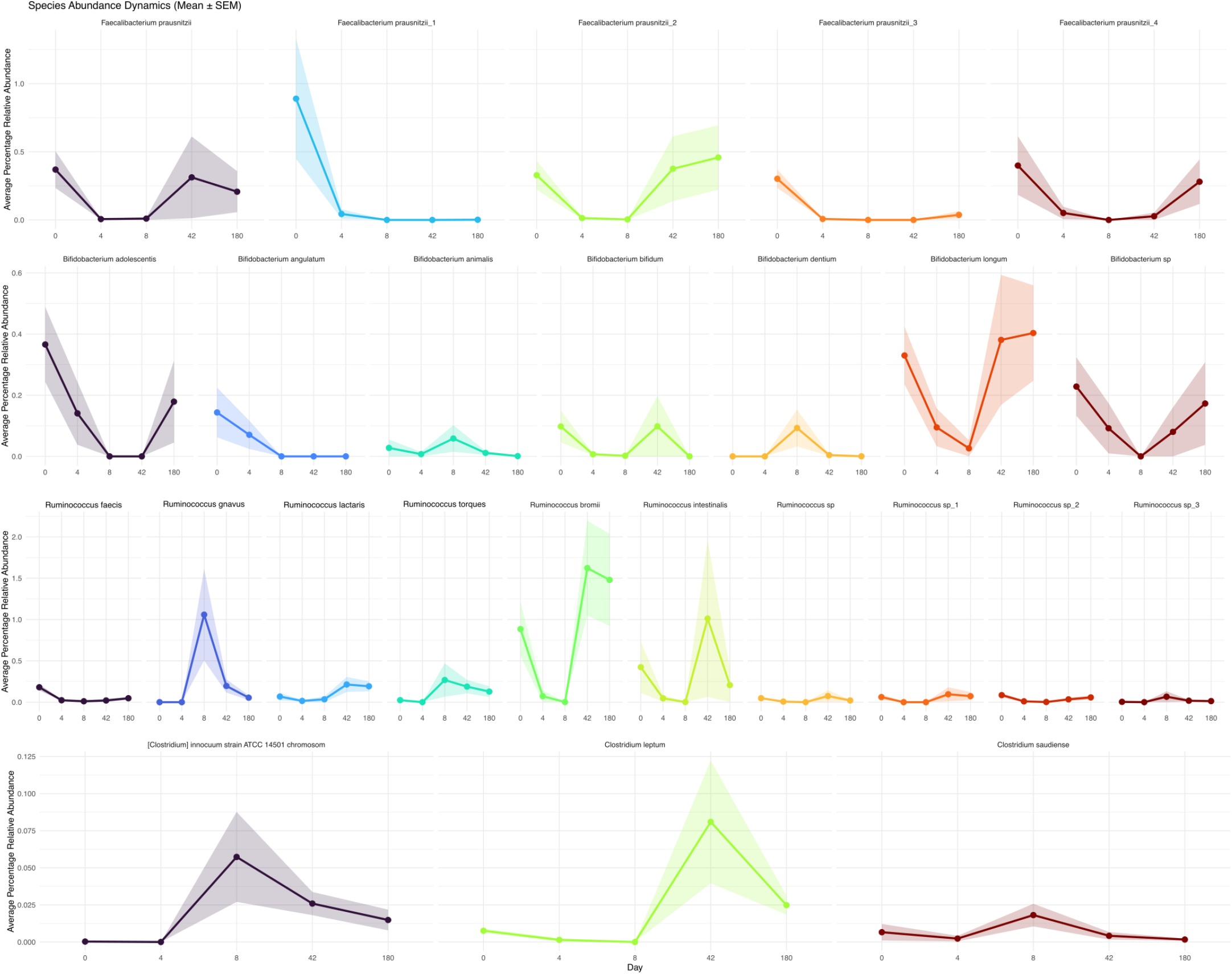
Strain-level relative abundance dynamics of representative taxa following antibiotic exposure. This figure extends manuscript Figure 2a by showing relative abundance trajectories across baseline, day 4, 8, 42 and 180 for selected taxa. As detected in the original study by Palleja et al. StrainSpy confirms that *Faecalibacterium prausnitzii* shows heterogeneous responses to the perturbation, with partial recovery of only some baseline-associated strains in some individuals at later time points, while other strains are lost from the population; this signal is detected in both cANI and abundance. In addition, *Bifidobacterium* spp. display a transient post-treatment shift in strain composition followed by gradual re-establishment of baseline-associated strains in a subset of individuals. *Ruminococcus* spp. exhibit similar strain turnover and partial recovery patterns across subjects. These trajectories are consistent with the broader patterns of community perturbation and recovery, highlighting strain-specific resilience, replacement, and persistence following antibiotic disturbance. However, such strain-level dynamics are not captured by species-level differential abundance analyses.

**Fig. S5.**
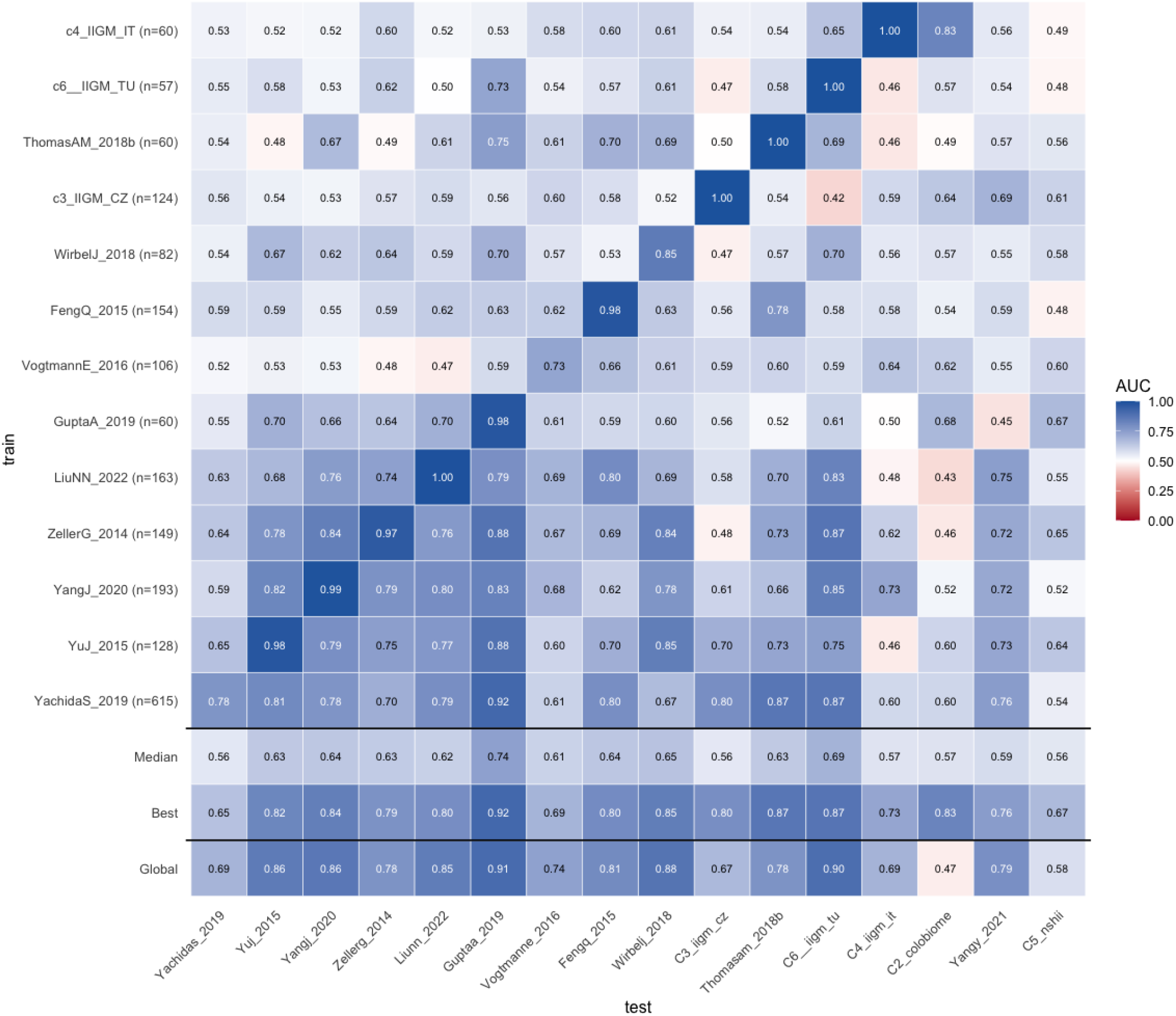
Evaluation of strain-level predictor portability across colorectal cancer microbiome cohorts. Heatmap showing pairwise cross-study prediction performance of elastic-net classifiers trained independently on each colorectal cancer microbiome cohort. Models were evaluated on all cohorts.

**Fig. S6.**
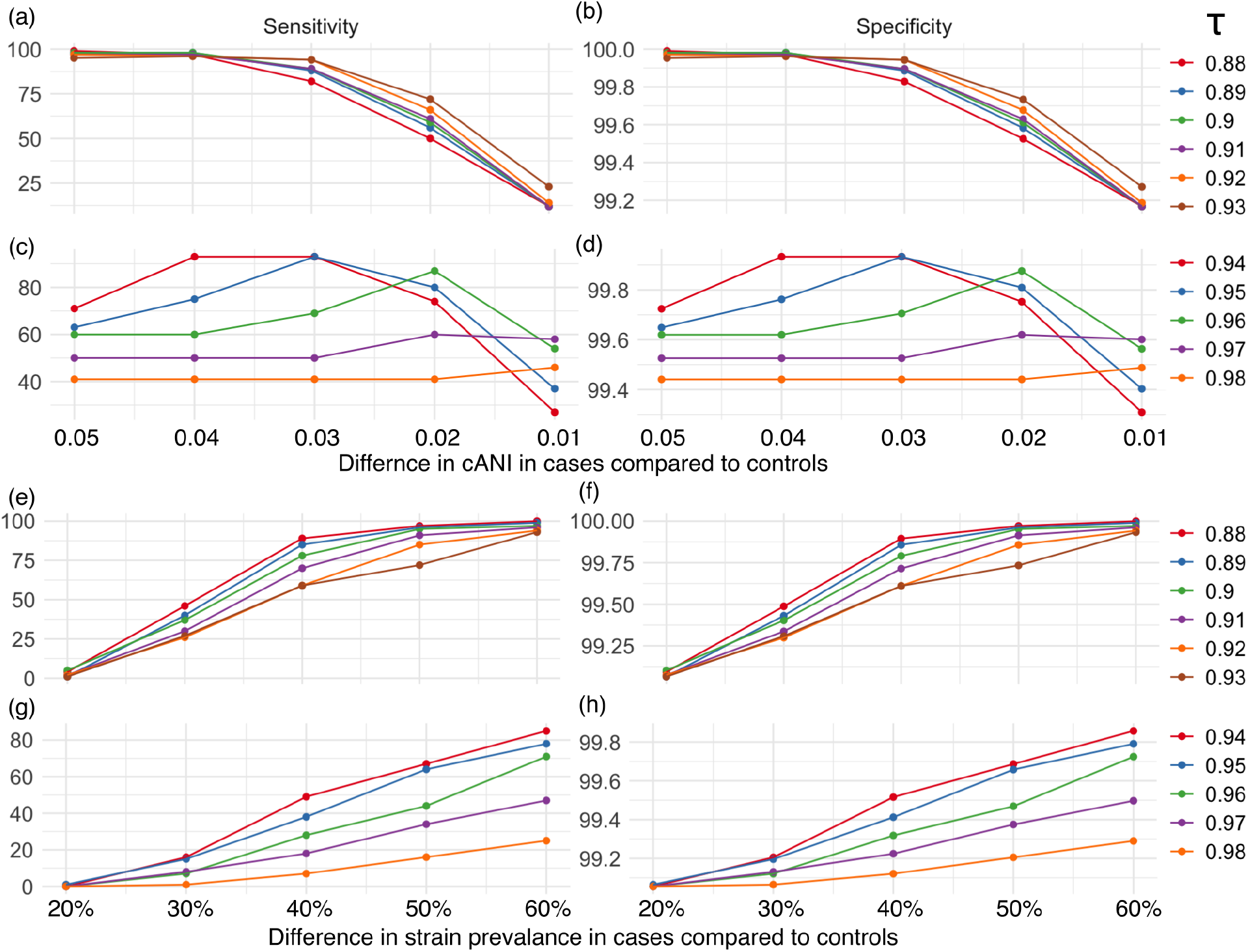
(a, c) Sensitivity and (b, d) specificity for simulations of strain identity differences between cases and controls. The x-axis indicates the simulated reduction in cANI between cases and controls, while line colour denotes the cANI threshold used to binarise strain presence for logistic regression. Panels (a) and (b) show thresholds (l≤ 0.93), for which performance is generally stable, whereas panels (c) and (d) show thresholds (≥ 0.94), where performance becomes increasingly variable and dependent on the chosen threshold. (e–h) Sensitivity and specificity for simulations of strain prevalence differences between cases and controls. The x-axis indicates the simulated proportion of depleted strains in the case group, with line colour again representing the cANI binarisation threshold. Similar threshold-dependent behaviour is observed, demonstrating that logistic regression performance is sensitive to the arbitrary threshold used to define strain presence and absence.

## References

1. Nicola K Petty, Nouri L Ben Zakour, Mitchell Stanton-Cook, Elizabeth Skippington, Makrina Totsika, Brian M Forde, Minh-Duy Phan, Danilo Gomes Moriel, Kate M Peters, Mark Davies, Benjamin A Rogers, Gordon Dougan, Jesús Rodriguez-Baño, Alvaro Pascual, Johann D D Pitout, Mathew Upton, David L Paterson, Timothy R Walsh, Mark A Schembri, and Scott A Beatson. Global dissemination of a multidrug resistant escherichia coli clone. Proc. Natl. Acad. Sci. U. S. A., 111(15):5694–5699, April 2014.

2. Cayetano Pleguezuelos-Manzano, Jens Puschhof, Axel Rosendahl Huber, Arne van Hoeck, Henry M Wood, Jason Nomburg, Carino Gurjao, Freek Manders, Guillaume Dalmasso, Paul B Stege, Fernanda L Paganelli, Maarten H Geurts, Joep Beumer, Tomohiro Mizutani, Yi Miao, Reinier van der Linden, Stefan van der Elst, Genomics England Research Consortium, K Christopher Garcia, Janetta Top, Rob J L Willems, Marios Giannakis, Richard Bonnet, Phil Quirke, Matthew Meyerson, Edwin Cuppen, Ruben van Boxtel, and Hans Clevers. Mutational signature in colorectal cancer caused by genotoxic pks+ e. coli. Nature, 580 (7802):269–273, April 2020.

3. Timothy J Dallman, Katri Jalava, Neville Q Verlander, David Gally, Claire Jenkins, Gauri Godbole, and Saheer Gharbia. Identification of domestic reservoirs and common exposures in an emerging lineage of shiga toxin-producing escherichia coli o157:h7 in england: a genomic epidemiological analysis. The Lancet Microbe, 3(8):e606–e615, 2022. ISSN 2666-5247. doi: 10.1016/S2666-5247(22)00089-1.

4. Danielle J Ingle, Marija Tauschek, David J Edwards, Dianna M Hocking, Derek J Pickard, Kristy I Azzopardi, Thakshila Amarasena, Vicki Bennett-Wood, Jaclyn S Pearson, Boubou Tamboura, et al. Evolution of atypical enteropathogenic e. coli by repeated acquisition of lee pathogenicity island variants. Nature Microbiology, 1(2):15010, 2016.

5. Marie Touchon, Amandine Perrin, Jorge Andre Moura De Sousa, Belinda Vangchhia, Samantha Burn, Claire L O’Brien, Erick Denamur, David Gordon, and Eduardo PC Rocha. Phylogenetic background and habitat drive the genetic diversification of escherichia coli. PLoS genetics, 16(6):e1008866, 2020.

6. Michael L Patnode, Janaki L Guruge, Juan J Castillo, Garret A Couture, Vincent Lombard, Nicolas Terrapon, Bernard Henrissat, Carlito B Lebrilla, and Jeffrey I Gordon. Strain-level functional variation in the human gut microbiota based on bacterial binding to artificial food particles. Cell Host Microbe, 29(4):664–673.e5, April 2021.

7. Vayu Maini Rekdal, Elizabeth N Bess, Jordan E Bisanz, Peter J Turnbaugh, and Emily P Balskus. Discovery and inhibition of an interspecies gut bacterial pathway for levodopa metabolism. Science, 364(6445):eaau6323, June 2019.

8. Se-Hoon Lee, Sung-Yup Cho, Youngmin Yoon, Changho Park, Jinyoung Sohn, Jin-Ju Jeong, Bu-Nam Jeon, Mongjoo Jang, Choa An, Suro Lee, Yun Yeon Kim, Gihyeon Kim, Sujeong Kim, Yunjae Kim, Gwang Bin Lee, Eun Ju Lee, Sang Gyun Kim, Hong Sook Kim, Yeongmin Kim, Hyun Kim, Hyun-Suk Yang, Sarang Kim, Seonggon Kim, Hayung Chung, Myeong Hee Moon, Myung Hee Nam, Jee Young Kwon, Sungho Won, Joon-Suk Park, George M Weinstock, Charles Lee, Kyoung Wan Yoon, and Hansoo Park. Bifidobacterium bifidum strains synergize with immune checkpoint inhibitors to reduce tumour burden in mice. Nat. Microbiol., 6(3):277–288, March 2021.

9. Andrew D Fernandes, Jennifer Ns Reid, Jean M Macklaim, Thomas A McMurrough, David R Edgell, and Gregory B Gloor. Unifying the analysis of high-throughput sequencing datasets: characterizing RNA-seq, 16S rRNA gene sequencing and selective growth experiments by compositional data analysis. Microbiome, 2(1):15, May 2014.

10. Huang Lin and Shyamal Das Peddada. Multigroup analysis of compositions of microbiomes with covariate adjustments and repeated measures. Nat. Methods, 21(1):83–91, January 2024.

11. William A Nickols, Thomas Kuntz, Jiaxian Shen, Sagun Maharjan, Himel Mallick, Eric A Franzosa, Kelsey N Thompson, Jacob T Nearing, and Curtis Huttenhower. MaAsLin 3:refining and extending generalized multivariable linear models for meta-omic association discovery. Nat. Methods, 23(3):554–564, March 2026.

12. Himel Mallick, Ali Rahnavard, Lauren J McIver, Siyuan Ma, Yancong Zhang, Long H Nguyen, Timothy L Tickle, George Weingart, Boyu Ren, Emma H Schwager, Suvo Chatterjee, Kelsey N Thompson, Jeremy E Wilkinson, Ayshwarya Subramanian, Yiren Lu, Levi Waldron, Joseph N Paulson, Eric A Franzosa, Hector Corrada Bravo, and Curtis Huttenhower. Multivariable association discovery in population-scale meta-omics studies. PLoS Comput. Biol., 17(11):e1009442, November 2021.

13. Jacob T Nearing, Gavin M Douglas, Molly G Hayes, Jocelyn MacDonald, Dhwani K Desai, Nicole Allward, Casey M A Jones, Robyn J Wright, Akhilesh S Dhanani, André M Comeau, and Morgan G I Langille. Microbiome differential abundance methods produce different results across 38 datasets. Nat. Commun., 13(1):342, January 2022.

14. Suguru Nishijima, Evelina Stankevic, Oliver Aasmets, Thomas S B Schmidt, Naoyoshi Nagata, Marisa Isabell Keller, Pamela Ferretti, Helene Bæk Juel, Anthony Fullam, Shahriyar Mahdi Robbani, Christian Schudoma, Johanne Kragh Hansen, Louise Aas Holm, Mads Israelsen, Robert Schierwagen, Nikolaj Torp, Anja Telzerow, Rajna Hercog, Stefanie Kandels, Diënty H M Hazenbrink, Manimozhiyan Arumugam, Flemming Bendtsen, Charlotte Brøns, Cilius Esmann Fonvig, Jens-Christian Holm, Trine Nielsen, Julie Steen Pedersen, Maja Sofie Thiele, Jonel Trebicka, Elin Org, Aleksander Krag, Torben Hansen, Michael Kuhn, Peer Bork, and GALAXY and MicrobLiver Consortia. Fecal microbial load is a major determinant of gut microbiome variation and a confounder for disease associations. Cell, 188(1):222–236.e15, January 2025.

15. Paul J McMurdie and Susan Holmes. Waste not, want not: why rarefying microbiome data is inadmissible. PLoS Comput. Biol., 10(4):e1003531, April 2014.

16. Jakob Wirbel, Morgan Essex, Sofia Kirke Forslund, and Georg Zeller. A realistic benchmark for differential abundance testing and confounder adjustment in human microbiome studies. Genome Biology, 25(1):247, September 2024.

17. Jim Shaw and Yun William Yu. Rapid species-level metagenome profiling and containment estimation with sylph. Nat Biotechnol, October 2024.

18. Luiz Irber, N Tessa Pierce-Ward, Mohamed Abuelanin, Harriet Alexander, Abhishek Anant, Keya Barve, Colton Baumler, Olga Botvinnik, Phillip Brooks, Daniel Dsouza, Laurent Gautier, Mahmudur Rahman Hera, Hannah Eve Houts, Lisa K Johnson, Fabian Klötzl, David Koslicki, Marisa Lim, Ricky Lim, Bradley Nelson, Ivan Ogasawara, Taylor Reiter, Camille Scott, Andreas Sjödin, Daniel Standage, S Joshua Swamidass, Connor Tiffany, Pranathi Vemuri, Erik Young, and C Titus Brown. sourmash v4: A multitool to quickly search, compare, and analyze genomic and metagenomic data sets. J. Open Source Softw., 9(98):6830, June 2024.

19. Brian D Ondov, Gabriel J Starrett, Anna Sappington, Aleksandra Kostic, Sergey Koren, Christopher B Buck, and Adam M Phillippy. Mash screen: high-throughput sequence containment estimation for genome discovery. Genome Biol., 20(1):232, November 2019.

20. Shaopeng Liu and David Koslicki. CMash: fast, multi-resolution estimation of k-mer-based jaccard and containment indices. Bioinformatics, 38(Suppl 1):i28–i35, June 2022.

21. Andrew R Ghazi, Kelsey N Thompson, Amrisha Bhosle, Zhendong Mei, Yan Yan, Fenglei Wang, Kai Wang, Eric A Franzosa, and Curtis Huttenhower. Quantifying metagenomic strain associations from microbiomes with anpan. January 2025.

22. John A Lees, Marco Galardini, Stephen D Bentley, Jeffrey N Weiser, and Jukka Corander. pyseer: a comprehensive tool for microbial pangenome-wide association studies. Bioinformatics, 34(24):4310–4312, December 2018.

23. Sarah G Earle, Chieh-Hsi Wu, Jane Charlesworth, Nicole Stoesser, N Claire Gordon, Timothy M Walker, Chris C A Spencer, Zamin Iqbal, David A Clifton, Katie L Hopkins, Neil Woodford, E Grace Smith, Nazir Ismail, Martin J Llewelyn, Tim E Peto, Derrick W Crook, Gil McVean, A Sarah Walker, and Daniel J Wilson. Identifying lineage effects when controlling for population structure improves power in bacterial association studies. Nat Microbiol, 1:16041, April 2016.

24. Gianmarco Piccinno, Kelsey N Thompson, Paolo Manghi, Andrew R Ghazi, Andrew Maltez Thomas, Aitor Blanco-Míguez, Francesco Asnicar, Katarina Mladenovic, Federica Pinto, Federica Armanini, Michal Punčochář, Elisa Piperni, Vitor Heidrich, Gloria Fackelmann, Giulio Ferrero, Sonia Tarallo, Long H Nguyen, Yan Yan, Nazim A Keles, Bilge G Tuna,Veronika Vymetalkova, Mario Trompetto, Vaclav Liska, Tomas Hucl, Pavel Vodicka, Beatrix Bencsiková, Martina Čarnogurská, Vlad Popovici, Federica Marmorino, Chiara Cremolini, Barbara Pardini, Francesca Cordero, Mingyang Song, Andrew T Chan, Lisa Derosa, Laurence Zitvogel, Curtis Huttenhower, Alessio Naccarati, Eva Budinska, and Nicola Segata. Pooled analysis of 3,741 stool metagenomes from 18 cohorts for cross-stage and strain-level reproducible microbial biomarkers of colorectal cancer. Nat. Med., 31(7):2416–2429, July 2025.

25. Albert Palleja, Kristian H Mikkelsen, Sofia K Forslund, Alireza Kashani, Kristine H Allin, Trine Nielsen, Tue H Hansen, Suisha Liang, Qiang Feng, Chenchen Zhang, Paul Theodor Pyl, Luis Pedro Coelho, Huanming Yang, Jian Wang, Athanasios Typas, Morten F Nielsen, Henrik Bjorn Nielsen, Peer Bork, Jun Wang, Tina Vilsbøll, Torben Hansen, Filip K Knop, Manimozhiyan Arumugam, and Oluf Pedersen. Recovery of gut microbiota of healthy adults following antibiotic exposure. Nat Microbiol, 3(11):1255–1265, November 2018.

26. Arthur E Frankel, Laura A Coughlin, Jiwoong Kim, Thomas W Froehlich, Yang Xie, Eugene P Frenkel, and Andrew Y Koh. Metagenomic shotgun sequencing and unbiased metabolomic profiling identify specific human gut microbiota and metabolites associated with immune checkpoint therapy efficacy in melanoma patients. Neoplasia, 19(10):848–855, October 2017.

27. V Gopalakrishnan, C N Spencer, L Nezi, A Reuben, M C Andrews, T V Karpinets, P A Prieto, D Vicente, K Hoffman, S C Wei, A P Cogdill, L Zhao, C W Hudgens, D S Hutchinson, T Manzo, M Petaccia de Macedo, T Cotechini, T Kumar, W S Chen, S M Reddy, R Szczepaniak Sloane, J Galloway-Pena, H Jiang, P L Chen, E J Shpall, K Rezvani, A M Alousi, R F Chemaly, S Shelburne, L M Vence, P C Okhuysen, V B Jensen, A G Swennes, F McAllister, E Marcelo Riquelme Sanchez, Y Zhang, E Le Chatelier, L Zitvogel, N Pons, J L Austin-Breneman, L E Haydu, E M Burton, J M Gardner, E Sirmans, J Hu, A J Lazar, T Tsujikawa, A Diab, H Tawbi, I C Glitza, W J Hwu, S P Patel, S E Woodman, R N Amaria, M A Davies, J E Gershenwald, P Hwu, J E Lee, J Zhang, L M Coussens, Z A Cooper, P A Futreal, C R Daniel, N J Ajami, J F Petrosino, M T Tetzlaff, P Sharma, J P Allison, R R Jenq, and J A Wargo. Gut microbiome modulates response to anti-PD-1 immunotherapy in melanoma patients. Science, 359(6371):97–103, January 2018.

28. Karla A Lee, Andrew Maltez Thomas, Laura A Bolte, Johannes R Björk, Laura Kist de Ruijter, Federica Armanini, Francesco Asnicar, Aitor Blanco-Miguez, Ruth Board, Neus Calbet-Llopart, Lisa Derosa, Nathalie Dhomen, Kelly Brooks, Mark Harland, Mark Harries, Emily R Leeming, Paul Lorigan, Paolo Manghi, Richard Marais, Julia Newton-Bishop, Luigi Nezi, Federica Pinto, Miriam Potrony, Susana Puig, Patricio Serra-Bellver, Heather M Shaw, Sabrina Tamburini, Sara Valpione, Amrita Vijay, Levi Waldron, Laurence Zitvogel, Moreno Zolfo, Elisabeth G E de Vries, Paul Nathan, Rudolf S N Fehrmann, Véronique Bataille, Geke A P Hospers, Tim D Spector, Rinse K Weersma, and Nicola Segata. Cross-cohort gut microbiome associations with immune checkpoint inhibitor response in advanced melanoma. Nat. Med., 28(3):535–544, March 2022.

29. Vyara Matson, Jessica Fessler, Riyue Bao, Tara Chongsuwat, Yuanyuan Zha, Maria-Luisa Alegre, Jason J Luke, and Thomas F Gajewski. The commensal microbiome is associated with anti-PD-1 efficacy in metastatic melanoma patients. Science, 359(6371):104–108, January 2018.

30. John A McCulloch, Diwakar Davar, Richard R Rodrigues, Jonathan H Badger, Jennifer R Fang, Alicia M Cole, Ascharya K Balaji, Marie Vetizou, Stephanie M Prescott, Miriam R Fernandes, Raquel G F Costa, Wuxing Yuan, Rosalba Salcedo, Erol Bahadiroglu, Soumen Roy, Richelle N DeBlasio, Robert M Morrison, Joe-Marc Chauvin, Quanquan Ding, Bochra Zidi, Ava Lowin, Saranya Chakka, Wentao Gao, Ornella Pagliano, Scarlett J Ernst, Amy Rose, Nolan K Newman, Andrey Morgun, Hassane M Zarour, Giorgio Trinchieri, and Amiran K Dzutsev. Intestinal microbiota signatures of clinical response and immune-related adverse events in melanoma patients treated with anti-PD-1. Nat. Med., 28(3):545–556, March 2022.

31. Christine N Spencer, Jennifer L McQuade, Vancheswaran Gopalakrishnan, John A Mc-Culloch, Marie Vetizou, Alexandria P Cogdill, Md A Wadud Khan, Xiaotao Zhang, Michael G White, Christine B Peterson, Matthew C Wong, Golnaz Morad, Theresa Rodgers, Jonathan H Badger, Beth A Helmink, Miles C Andrews, Richard R Rodrigues, Andrey Morgun, Young S Kim, Jason Roszik, Kristi L Hoffman, Jiali Zheng, Yifan Zhou, Yusra B Medik, Laura M Kahn, Sarah Johnson, Courtney W Hudgens, Khalida Wani, Pierre-Olivier Gaudreau, Angela L Harris, Mohamed A Jamal, Erez N Baruch, Eva Perez-Guijarro, Chi-Ping Day, Glenn Merlino, Barbara Pazdrak, Brooke S Lochmann, Robert A Szczepaniak-Sloane, Reetakshi Arora, Jaime Anderson, Chrystia M Zobniw, Eliza Posada, Elizabeth Sirmans, Julie Simon, Lauren E Haydu, Elizabeth M Burton, Linghua Wang, Minghao Dang, Karen Clise-Dwyer, Sarah Schneider, Thomas Chapman, Nana-Ama A S Anang, Sheila Duncan, Joseph Toker, Jared C Malke, Isabella C Glitza, Rodabe N Amaria, Hussein A Tawbi, Adi Diab, Michael K Wong, Sapna P Patel, Scott E Woodman, Michael A Davies, Merrick I Ross, Jeffrey E Gershenwald, Jeffrey E Lee, Patrick Hwu, Vanessa Jensen, Yardena Samuels, Ravid Straussman, Nadim J Ajami, Kelly C Nelson, Luigi Nezi, Joseph F Petrosino, P Andrew Futreal, Alexander J Lazar, Jianhua Hu, Robert R Jenq, Michael T Tetzlaff, Yan Yan, Wendy S Garrett, Curtis Huttenhower, Padmanee Sharma, Stephanie S Watowich, James P Allison, Lorenzo Cohen, Giorgio Trinchieri, Carrie R Daniel, and Jennifer A Wargo. Dietary fiber and probiotics influence the gut microbiome and melanoma immunotherapy response. Science, 374(6575):1632–1640, December 2021.

32. Ashray Gunjur, Yan Shao, Timothy Rozday, Oliver Klein, Andre Mu, Bastiaan W Haak, Ben Markman, Damien Kee, Matteo S Carlino, Craig Underhill, Sophia Frentzas, Michael Michael, Bo Gao, Jodie Palmer, Jonathan Cebon, Andreas Behren, David J Adams, and Trevor D Lawley. A gut microbial signature for combination immune checkpoint blockade across cancer types. Nat Med, 30(3):797–809, March 2024.

33. Daniel J Wilson. The harmonic mean p-value for combining dependent tests. Proc. Natl. Acad. Sci. U. S. A., 116(4):1195–1200, January 2019.

34. Paolo Manghi, Giacomo Antonello, Lucas Schiffer, Davide Golzato, Andres Wokaty, Francesco Beghini, Chloe Mirzayi, Kaelyn Long, Kai Gravel-Pucillo, Gianmarco Piccinno, Samuel David Gamboa-Tuz, Arianna Bonetti, Giacomo D’Amato, Rimsha Azhar, Kelly Eckenrode, Fatima Zohra, Valentina Giunchiglia, Marisa Keller, Anna Pedrotti, Ilya Likhotkin, Shaimaa Elsafoury, Ludwig Geistlinger, Aitor Blanco-Miguez, Andrew Maltez Thomas, Moreno Zolfo, Marcel Ramos, Mireia Valles-Colomer, Sabrina Tamburini, Francesco As-nicar, Heidi E Jones, Curtis Huttenhower, Vincent Carey, Sean Davis, Edoardo Pasolli, Sehyun Oh, Nicola Segata, and Levi Waldron. Meta-analysis of 22,710 human microbiome metagenomes defines an oral-to-gut microbial enrichment score and associations with host health and disease. Nat. Commun., 17(1):196, December 2025.

35. David Zeevi, Tal Korem, Niv Zmora, David Israeli, Daphna Rothschild, Adina Weinberger, Orly Ben-Yacov, Dar Lador, Tali Avnit-Sagi, Maya Lotan-Pompan, Jotham Suez, Jemal Ali Mahdi, Elad Matot, Gal Malka, Noa Kosower, Michal Rein, Gili Zilberman-Schapira, Lenka Dohnalová, Meirav Pevsner-Fischer, Rony Bikovsky, Zamir Halpern, Eran Elinav, and Eran Segal. Personalized nutrition by prediction of glycemic responses. Cell, 163(5):1079–1094, November 2015.

36. Robert Kubinec. Ordered beta regression: A parsimonious, Well-Fitting model for continuous data with lower and upper bounds. Political Analysis, 31(4):519–536, October 2023.

37. Zhejian Yu. Art_modern: An accelerated ART simulator of diverse next-generation sequencing reads. February 2026.

38. Weichun Huang, Leping Li, Jason R Myers, and Gabor T Marth. ART: a next-generation sequencing read simulator. Bioinformatics, 28(4):593–594, February 2012.

39. Duy Tin Truong, Adrian Tett, Edoardo Pasolli, Curtis Huttenhower, and Nicola Segata. Microbial strain-level population structure and genetic diversity from metagenomes. Genome Res., 27(4):626–638, April 2017.

40. Fariba Akrami, Hossein Jamali, Mansoor Kodori, and Charles M. Dozois. Papb family regulators as master switches of fimbrial expression. Microorganisms, 13(8), 2025. ISSN 2076-2607. doi: 10.3390/microorganisms13081939.

41. William R Schwan. Regulation of fim genes in uropathogenic escherichia coli. World journal of clinical infectious diseases, 1(1):17, 2011.

42. Louise Roer, Veronika Tchesnokova, Rosa Allesøe, Mariya Muradova, Sujay Chattopadhyay, Johanne Ahrenfeldt, Martin CF Thomsen, Ole Lund, Frank Hansen, Anette M Hammerum, et al. Development of a web tool for escherichia coli subtyping based on fimh alleles. Journal of clinical microbiology, 55(8):2538–2543, 2017.

43. Hiroshi Ogasawara, Toshiyuki Ishizuka, Shuhei Hotta, Michiko Aoki, Tomohiro Shimada, and Akira Ishihama. Novel regulators of the csgd gene encoding the master regulator of biofilm formation in escherichia coli k-12. Microbiology, 166(9):880–890, 2020.

44. Martha Zepeda-Rivera, Samuel S Minot, Heather Bouzek, Hanrui Wu, Aitor Blanco-Míguez, Paolo Manghi, Dakota S Jones, Kaitlyn D LaCourse, Ying Wu, Elsa F McMahon, Soon-Nang Park, Yun K Lim, Andrew G Kempchinsky, Amy D Willis, Sean L Cotton, Susan C Yost, Ewa Sicinska, Joong-Ki Kook, Floyd E Dewhirst, Nicola Segata, Susan Bullman, and Christopher D Johnston. A distinct fusobacterium nucleatum clade dominates the colorectal cancer niche. Nature, 628(8007):424–432, April 2024.

45. Marie Vétizou, Jonathan M Pitt, Romain Daillère, Patricia Lepage, Nadine Waldschmitt, Caroline Flament, Sylvie Rusakiewicz, Bertrand Routy, Maria P Roberti, Connie P M Duong, Vichnou Poirier-Colame, Antoine Roux, Sonia Becharef, Silvia Formenti, Encouse Golden, Sascha Cording, Gerard Eberl, Andreas Schlitzer, Florent Ginhoux, Sridhar Mani, Takahiro Yamazaki, Nicolas Jacquelot, David P Enot, Marion Bérard, Jérôme Nigou, Paule Opolon, Alexander Eggermont, Paul-Louis Woerther, Elisabeth Chachaty, Nathalie Chaput, Caroline Robert, Christina Mateus, Guido Kroemer, Didier Raoult, Ivo Gomperts Boneca, Franck Carbonnel, Mathias Chamaillard, and Laurence Zitvogel. Anticancer immunotherapy by CTLA-4 blockade relies on the gut microbiota. Science, 350(6264):1079–1084, November 2015.

46. Maik Luu, Zeno Riester, Adrian Baldrich, Nicole Reichardt, Samantha Yuille, Alessandro Busetti, Matthias Klein, Anne Wempe, Hanna Leister, Hartmann Raifer, Felix Picard, Khalid Muhammad, Kim Ohl, Rossana Romero, Florence Fischer, Christian A Bauer, Magdalena Huber, Thomas M Gress, Matthias Lauth, Sophia Danhof, Tobias Bopp, Thomas Nerreter, Imke E Mulder, Ulrich Steinhoff, Michael Hudecek, and Alexander Visekruna. Microbial short-chain fatty acids modulate CD8+ T cell responses and improve adoptive immunotherapy for cancer. Nat. Commun., 12(1):4077, July 2021.

47. Annabell Bachem, Michele Clarke, Geraldine Kong, Ilaryia Tarasova, Lachlan Dryburgh, Lindsay Kosack, Ariane R Lee, Lewis D Newland, Teagan Wagner, Emma Bawden, Kah Min Yap, Michael D Wilson, Sven Engel, Kayla R Wilson, Brendan E Russ, Kshitij Tandon, Peter Kar Han Lau, Grant McArthur, Vanessa R Marcelino, Paul A Beavis, Stephen J Turner, Jason Waithman, Timothy P Stinear, Shahneen Sandhu, Jan Schröder, Thomas Gebhardt, and Sammy Bedoui. Microbiota-derived butyrate promotes a FOXO1-induced stemness program and preserves CD8+ T cell immunity against melanoma. Immunity, 58(11):2799–2813.e8, November 2025.

48. Lysanne Desharnais, Anikka Swaby, Meriem Messaoudene, Samuel Doré, Miranda W Yu, Benoit Fiset, Valérie Breton, Mayra Ponce, Yongjia Hu, Liam Wilson, Mark Sorin, Ye Wang, Ken Dewar, Michael Pollak, Arielle Elkrief, Bertrand Routy, Logan A Walsh, and Daniela F Quail. Diet–microbiome synergy underlies obesity-associated immunotherapy efficacy. Nature, July 2026.

49. Gary D Wu, Jun Chen, Christian Hoffmann, Kyle Bittinger, Ying-Yu Chen, Sue A Keilbaugh, Meenakshi Bewtra, Dan Knights, William A Walters, Rob Knight, Rohini Sinha, Erin Gilroy, Kernika Gupta, Robert Baldassano, Lisa Nessel, Hongzhe Li, Frederic D Bushman, and James D Lewis. Linking long-term dietary patterns with gut microbial enterotypes. Science, 334(6052):105–108, October 2011.

50. Lawrence A David, Corinne F Maurice, Rachel N Carmody, David B Gootenberg, Julie E Button, Benjamin E Wolfe, Alisha V Ling, A Sloan Devlin, Yug Varma, Michael A Fischbach, Sudha B Biddinger, Rachel J Dutton, and Peter J Turnbaugh. Diet rapidly and reproducibly alters the human gut microbiome. Nature, 505(7484):559–563, January 2014.

51. Martin Morgan, Valerie Obenchain, Jim Hester, Hervé Pagès. SummarizedExperiment, 2017.

52. Donovan H Parks, Pierre-Alain Chaumeil, Aaron J Mussig, Christian Rinke, Maria Chuvochina, and Philip Hugenholtz. GTDB release 10: a complete and systematic taxonomy for 715 230 bacterial and 17 245 archaeal genomes. Nucleic Acids Res., 54(D1):D743–D754, January 2026.

53. Aitor Blanco-Míguez, Francesco Beghini, Fabio Cumbo, Lauren J McIver, Kelsey N Thompson, Moreno Zolfo, Paolo Manghi, Leonard Dubois, Kun D Huang, Andrew Maltez Thomas, William A Nickols, Gianmarco Piccinno, Elisa Piperni, Michal Punčochář, Mireia Valles-Colomer, Adrian Tett, Francesca Giordano, Richard Davies, Jonathan Wolf, Sarah E Berry,Tim D Spector, Eric A Franzosa, Edoardo Pasolli, Francesco Asnicar, Curtis Huttenhower, and Nicola Segata. Extending and improving metagenomic taxonomic profiling with uncharacterized species using MetaPhlAn 4. Nat Biotechnol, 41(11):1633–1644, November 2023.

54. Maeve McGillycuddy, David I Warton, Gordana Popovic, and Benjamin M Bolker. Parsimoniously fitting large multivariate random effects in glmmTMB. J. Stat. Softw., 112(1), 2025.

55. Mollie E Brooks, Kasper Kristensen, Koen J van Benthem, Arni Magnusson, Casper W Berg, Anders Nielsen, Hans J Skaug, Martin Mächler, and Benjamin M Bolker. glmmTMB balances speed and flexibility among packages for zero-inflated generalized linear mixed modeling. The R Journal, 9(2):378–400, December 2017.

56. Raydonal Ospina and Silvia L.P. Ferrari. A general class of zero-or-one inflated beta regression models. Computational Statistics and Data Analysis, 56(6):1609–1623, 2012. ISSN 0167-9473. doi: 10.1016/j.csda.2011.10.005.

57. Jared Huling. fastglm: Fast and stable fitting of generalized linear models using ‘RcppEigen’, March 2019. Title of the publication associated with this dataset: CRAN: Contributed Packages.

58. Francisco Cribari-Neto and Achim Zeileis. Beta regression inr. J. Stat. Softw., 34(2), 2010.

59. Jim Shaw and Yun William Yu. Fast and robust metagenomic sequence comparison through sparse chaining with skani. Nat. Methods, 20(11):1661–1665, November 2023.

60. Katarzyna Sidorczuk, Michał Burdukiewicz, Klara Cerk, Joachim Fritscher, Robert A Kingsley, Peter Schierack, Falk Hildebrand, and Rafał Kolenda. adhesiomer: a tool for escherichia coli adhesin classification and analysis. BMC Genomics, 25(1):609, June 2024.

61. Martin Hunt, Alison E Mather, Leonor Sánchez-Busó, Andrew J Page, Julian Parkhill, Jacqueline A Keane, and Simon R Harris. ARIBA: rapid antimicrobial resistance genotyping directly from sequencing reads. Microb. Genom., 3(10):e000131, October 2017.

62. Douglas Bates, Martin Mächler, Ben Bolker, and Steve Walker. Fitting linear Mixed-Effects models using lme4. June 2014.

63. Anna Ly, Rune Haubo Bojesen Christensen, Douglas Bates, Martin Maechler, and Benjamin M Bolker. Fitting generalized linear Mixed-Effects models using lme4. July 2026.

64. Max Kuhn. Building predictive models inRUsing thecaretpackage. J. Stat. Softw., 28(5), 2008.

65. Jerome Friedman, Trevor Hastie, and Rob Tibshirani. Regularization paths for generalized linear models via coordinate descent. J. Stat. Softw., 33(1):1–22, 2010.

## Supplementary References

1. Andreas Groll and Gerhard Tutz. Variable selection for generalized linear mixed models by l 1-penalized estimation. Statistics and Computing, 24(2):137–154, 2014.

2. Mollie E Brooks, Kasper Kristensen, Koen J Van Benthem, Arni Magnusson, Casper W Berg, Anders Nielsen, Hans J Skaug, Martin Mächler, and Benjamin M Bolker. glmmtmb balances speed and flexibility among packages for zero-inflated generalized linear mixed modeling. 2017.

3. Robert Kubinec. Ordered beta regression: a parsimonious, well-fitting model for continuous data with lower and upper bounds. Political analysis, 31(4):519–536, 2023.

4. Jim Shaw and Yun William Yu. Rapid species-level metagenome profiling and containment estimation with sylph. Nature Biotechnology, 43(8):1348–1359, 2025.

